# Synergistic neural integration is greater downstream of recurrent information flow in organotypic cortical cultures

**DOI:** 10.1101/2020.05.12.091215

**Authors:** Samantha P. Sherrill, Nicholas M. Timme, John M. Beggs, Ehren L. Newman

## Abstract

The directionality of network information flow dictates how networks process information. A central component of information processing in both biological and artificial neural networks is their ability to perform synergistic integration–a type of computation. We established previously that synergistic integration varies directly with the strength of feedforward information flow. However, the relationships between both recurrent and feedback information flow and synergistic integration remain unknown. To address this, we analyzed the spiking activity of hundreds of neurons in organotypic cultures of mouse cortex. We asked how empirically observed synergistic integration varied with local functional network structure that was categorized into motifs with varying recurrent and feedback information flow. We found that synergistic integration was elevated in motifs with greater recurrent information flow beyond that expected from the local feedforward information flow. Feedback information flow was interrelated with feedforward information flow and was associated with decreased synergistic integration. Our results indicate that synergistic integration is distinctly influenced by the directionality of local information flow.

**Author Summary:** Networks compute information. That is, they modify inputs to generate distinct outputs. These computations are an important component of network information processing. Knowing how the routing of information in a network influences computation is therefore crucial. Here we asked how a key form of computation—synergistic integration—is related to the direction of local information flow in networks of spiking cortical neurons. Specifically, we asked how information flow between input neurons (i.e., recurrent information flow) and information flow from output neurons to input neurons (i.e., feedback information flow) was related to the amount of synergistic integration performed by output neurons. We found that greater synergistic integration occurred where there was more recurrent information flow. And, lesser synergistic integration occurred where there was more feedback information flow relative to feedforward information flow. These results show that computation, in the form of synergistic integration, is distinctly influenced by the directionality of local information flow. Such work is valuable for predicting where and how network computation occurs and for designing networks with desired computational abilities.

## Introduction

Feedforward, recurrent and feedback connections are important for information processing in both artificial and biological neural networks [1–5]. Whether these connections represent the strength of a synapse, or the amount of information transmission between two nodes (i.e. information flow), the directionality of these connections–feedforward, recurrent (lateral) or feedback–influences how the network processes information. A component of information processing that is central to both biological and artificial neural networks is their ability to perform synergistic integration, a form of computation. Feedforward information flow has been previously shown to be a strong predictor of synergistic integration in biological cortical circuits [6]. However, the influence of recurrent and feedback information flow on synergistic integration is unclear. Understanding how each of these directed functional connections influences the computational properties of neural networks is a critical step in understanding how neural networks compute. Here, we examine this in the context of cortical networks, using a motif-style, information theoretic analysis of high-density *in vitro* recordings of spiking neurons.

Synergistic integration refers to the synergistic combination of existing information to derive new information which is “greater than the sum of the parts.” Thus, it is a proxy for a form of non-trivial computation. Synergistic integration can be measured as the synergy [7] that emerges when a given neuron integrates input from two other neurons [8,9]. This approach has been used effectively before [6,8,10,11]. Here, we leveraged this approach to determine how recurrent and feedback information flow relate to synergistic integration.

Recurrence—whether physical in the form of neuronal processes or functional in the form of information flow—is believed to be important for higher order functions including memory processes (e.g. recollection, recognition). This is due to their generation of attractor-like, pattern completion activity [12–20]. This type of activity involves the combination of diverse features to form representations, also contributing to the interpretation and categorization of representations [21–27]. These studies also show that greater interpretability of images and object categories occurs at latencies beyond those of known feedforward processes. Relatedly, in artificial neural networks, recurrent connections serve to expand computational power by extending operations in time, requiring a smaller network to carry out the same operations as a larger, purely feedforward network [3,28]. Thus, controlling for the size of a network, the use of recurrence can improve network operation.

Feedback—whether physical or functional, local or long-range—is believed to implement top-down, goal-driven attention and perception, which involves the preferential activation of lower level neurons by higher level neurons [e.g., 29-31; for reviews, see 32,33]. Due to its topdown nature, feedback also plays a role in the gating and rerouting of information flow, as well as error prediction [for related reviews see 34-38]. Relatedly, feedback is associated with increased surround suppression, reducing the range of stimuli to which lower-level neurons respond [39,40]. From this perspective, feedback reduces the variance with which lower-level neurons can account for variance in higher-level neurons.

Here, we tested how recurrence and feedback relate to synergistic integration in functional networks observed in cortex (Figure 1). Across motifs, we found that recurrence was positively related to synergistic integration. Feedback, conversely, was negatively related to synergistic integration, but this negative relationship was accounted for by concurrent shifts in feedforward information flow.

**Fig 1.**
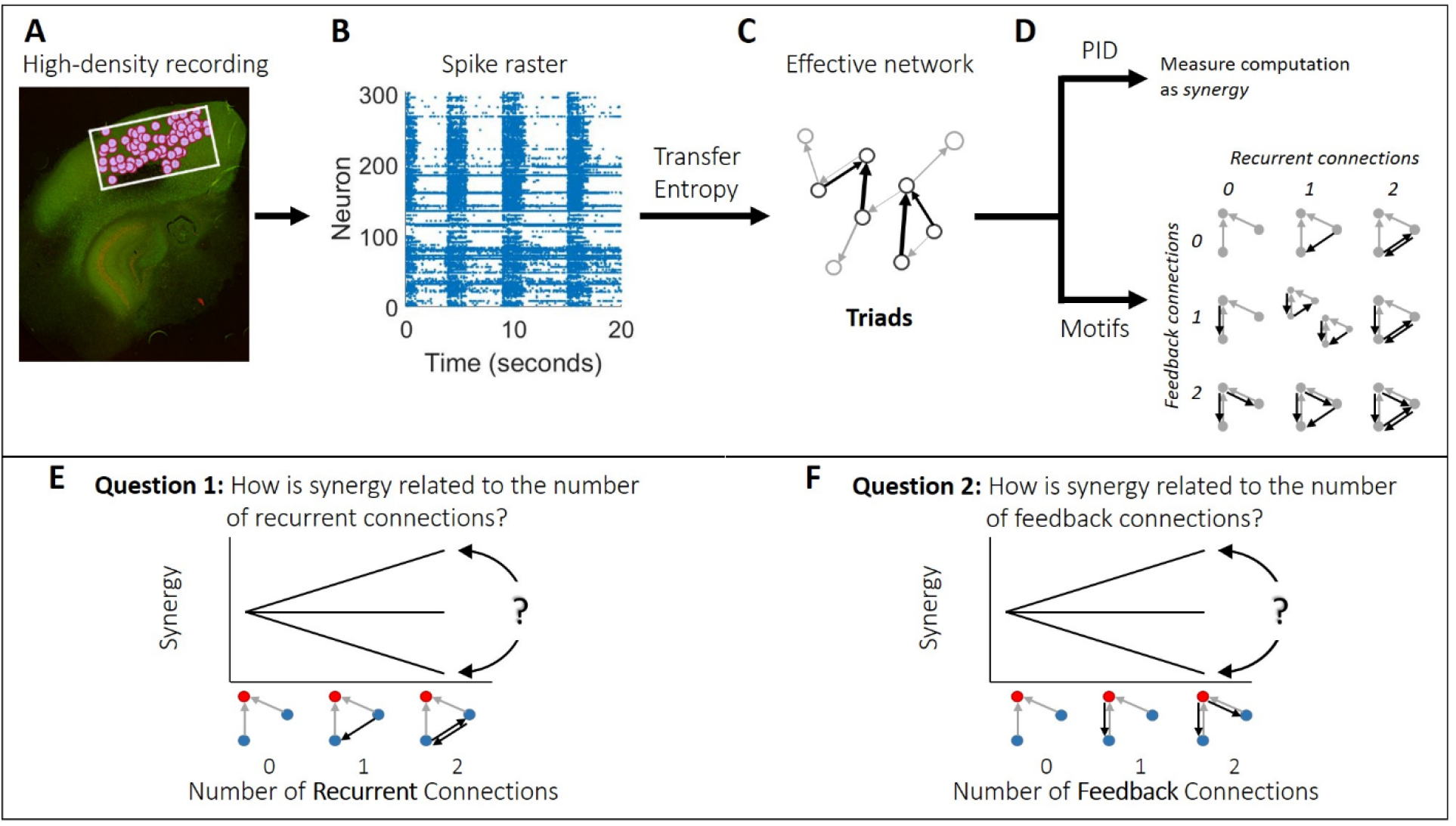
Methodological approach taken to ask how synergy is related to recurrent and feedback information flow in organotypic cultures of mouse cortex. (A) Hour-long recordings of spiking activity were collected *in vitro* from organotypic cultures of mouse somatosensory cortex using a high density 512-channel multielectrode array. (B) Spike sorting yielded spike trains of hundreds of well-isolated individual neurons per recording. (C) Transfer entropy was used to identify significant information flow between neuron pairs. This comprised the effective connections (edges) in a functional network. The resulting effective networks were analyzed to identify all triads consisting of two edges connecting to a common receiver. (D) For each triad, we quantified the amount of synergy via partial information decomposition (PID). We also identified all possible relevant motifs and arranged them according to the number of recurrent and feedback edges they contained. (E-F) We sought to answer two questions. (E) Is synergy positively related, negatively related, or unrelated to the number of recurrent edges? (F) Is synergy positively related, negatively related, or unrelated to the number of feedback edges? Triads consist of two transmitter neurons (blue), each with feedforward edges (gray arrows) connecting to a receiver neuron (red). Black arrows depict recurrent and feedback edges on the left and right, respectively.

## Results

We asked how the number of recurrent and feedback edges in triadic motifs was related to the amount of synergistic integration by analyzing hour long recordings of spiking activity from organotypic cultures of mouse somatosensory cortex (n = 25), as summarized in Figure 1. Recordings yielded between 98 and 594 well-isolated neurons (mean = 309). The average firing rate among neurons was 2.1 Hz [2.0 Hz, 2.2 Hz] and neurons exhibited rhythmic bursts of activity (Fig 1B) as characterized previously [41–42]. We identified effective connections between neurons in each recording as those that had significant transfer entropy such that the observed value was greater than 99.9% of values obtained from a jittering procedure (i.e. p<0.001). We then identified all synergistic 3-node motifs. Synergistic motifs were those which included two transmitter nodes sending inputs to the same receiver-node. Motifs without this structure were excluded because we were only concerned with the motifs’ ability to perform synergistic integration. The set of motifs included in our analyses are shown in Figure 2. We quantified the amount of synergistic integration performed by the receiver based on its inputs using ‘synergy,’ a term derived from partial information decomposition. Consistent with previous work [6, 8, 10], synergy was normalized to reflect the proportion of the receiving neuron entropy for which it accounted and to control for variable entropy across triads and networks. Across triads, we asked whether normalized synergy was positively or negatively related to the number of recurrent and feedback edges in the corresponding motif. This analysis was repeated at three timescales relevant to the delay of synaptic transmission, which has been reported to span 1-20 ms [43–44]. The three timescales (0.05-3 ms, 1.6-6.4 ms, and 3.5-14 ms), covering a range of 0.05-14 ms, were determined by the granularity of the data binning and the delay between bins. More detail regarding the structure of the timescales is given in the Supporting Information. All summary statistics are reported as medians or means, as indicated, followed by 95% bootstrap confidence intervals in brackets.

**Fig 2.**
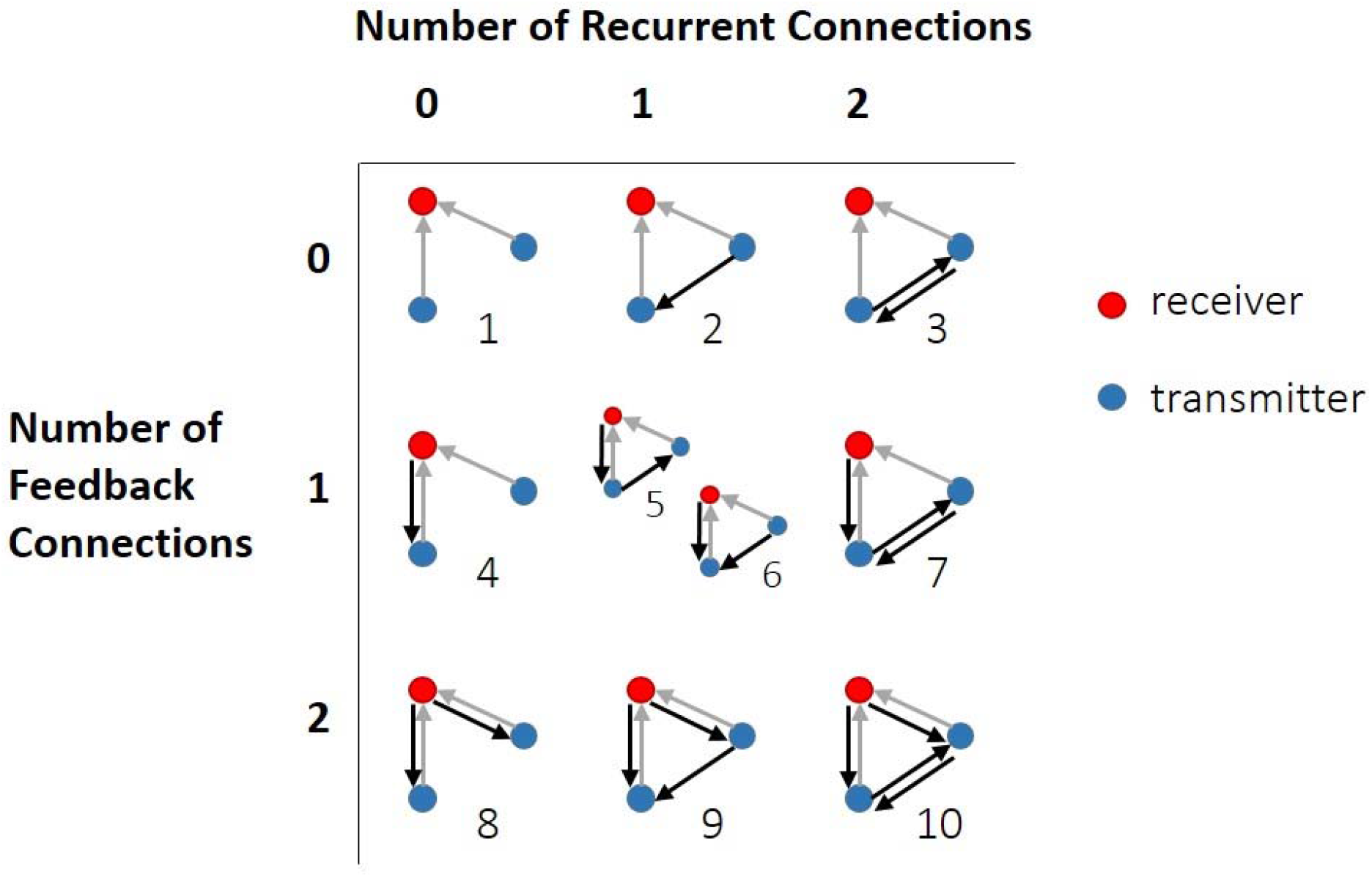
Set of synergistic 3-node motifs. Synergistic motifs were those in which both transmitters (blue dots) sent input (gray arrows) to the same receiver (red dots). Motifs were arranged in order (1-10) of the number of feedback and recurrent edges they contained (black arrows); either 0, 1 or 2.

### Recurrence predicts increased normalized synergy, feedback predicts decreased normalized synergy

To examine the relationships between normalized synergy and the number of both recurrent and feedback effective connections, we used a motif-style analysis [45]. We first quantified the mean normalized synergy for each of the 10 synergistic motifs in every network (Figure 3). We then compared the mean normalized synergy in recurrent motifs (those with more recurrent than feedback edges) to the mean normalized synergy in feedback motifs (those with more feedback than recurrent edges). We observed significantly greater normalized synergy in recurrent motifs (Fig 3, orange) compared to feedback motifs (Fig 3, green) (mean = 0.011 vs. 0.007, z_s.r_ = 6.31, n=75, p<1×10^−9^). To determine how recurrent and feedback edges affect normalized synergy relative to baseline levels, we compared the observed synergy in each motif to the normalized synergy in the default motif (with 0 recurrent and 0 feedback edges; Fig 3). We found that triads with recurrent motifs had significantly greater normalized synergy than those with the default motif (z_s.r_ = 5.80, n=75, p<1×10^−8^). Conversely, triads with feedback motifs had significantly less normalized synergy than those with the default motif (z_s.r_ = −2.61, n=75, p=0.009). In addition to the comparison to baseline normalized synergy, we compared the observed synergy values to those observed when motif labels were randomly permuted across triads. We observed the same qualitative pattern of results here as in the comparison to baseline synergy levels. We found that recurrent motifs had significantly greater normalized synergy than expected by chance (z_s.r_ = 5.93, n=75, p<1×10^−8^). Conversely, feedback motifs had significantly less normalized synergy than expected by chance (z_s.r_ = −3.96, n=75, p<1×10^−4^).

**Fig 3.**
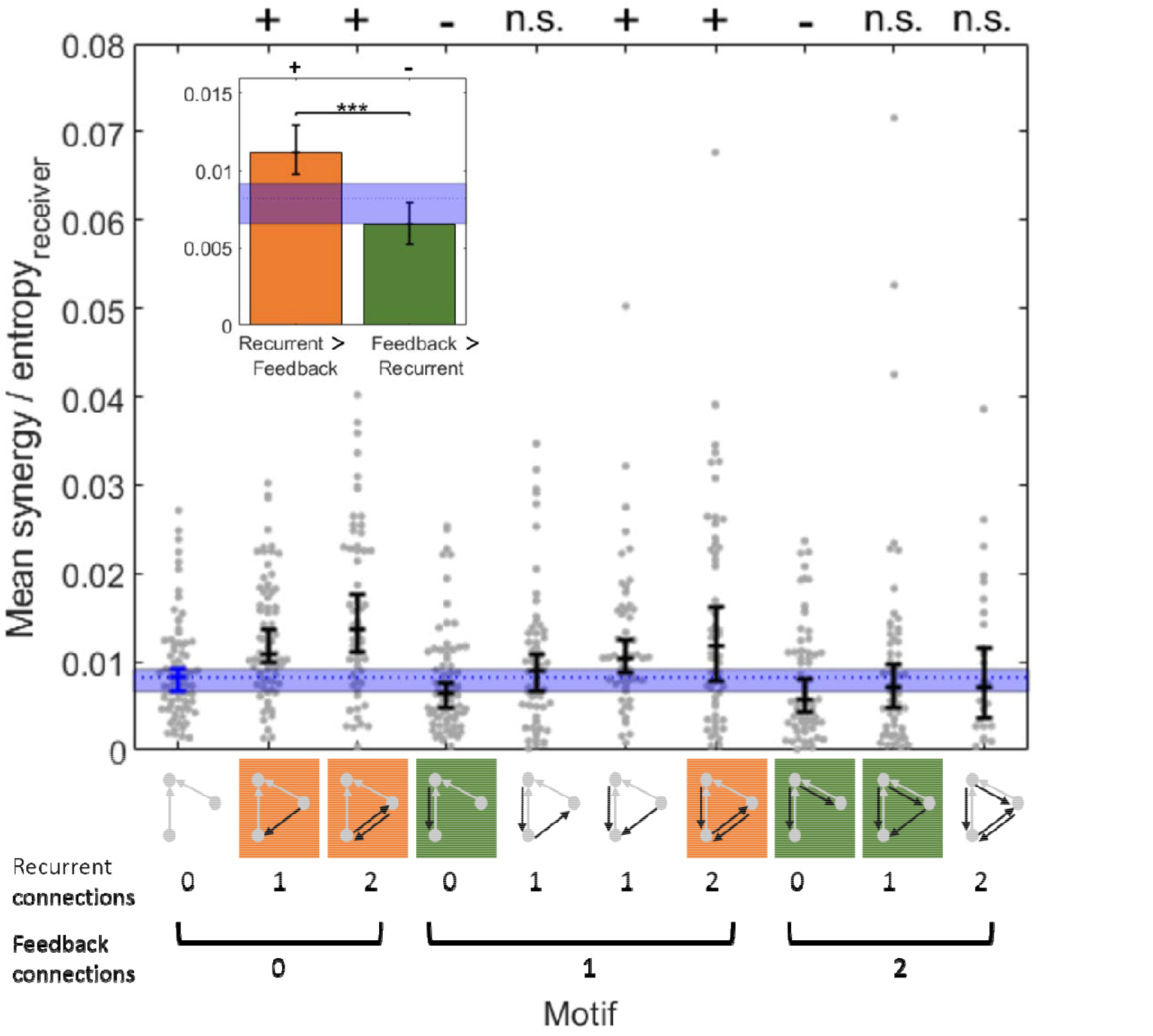
Normalized synergy was greater in recurrent motifs than in feedback motifs. Point clouds show the mean synergy value for each of the 75 networks analyzed for each type of motif. For distributions in which not all networks exhibited the motif, n<75. Central tendency and error bars depict the median and the 95% bootstrap confidence interval around the median. The motifs are graphically depicted below the x-axis and are organized by the number of feedback and recurrent edges. Motifs with more recurrent than feedback edges are indicated in orange. Motifs with more feedback than recurrent edges are indicated in green. Inset: Synergy values from motifs with greater recurrence or greater feedback were aggregated to directly compare the mean synergy. In both panels, the median (dotted line) and 95% bootstrap confidence interval (blue region) for baseline synergy values in default motifs (with 0 recurrent and 0 feedback edges) is shown. Significance indicators: ‘+’ and ‘-’ indicates p<0.01 by a two-tailed test wherein ‘+’ indicates significantly more than baseline and ‘-’ indicates significantly less than baseline; *** p<1×10^−9^.

To assess how normalized synergy varies across the multiple levels of recurrence and feedback, we grouped the 10 synergistic motifs into 9 categories based on the number of feedback and recurrent edges they contained (Fig 4A). A two-factor (recurrent and feedback), repeated measures ANOVA, with three levels of each factor (0,1 or 2 edges), was conducted to examine the main effects of recurrence and feedback on synergy and to test for an interaction effect. The main effect of recurrence was significant (*F*(2,148)=7, p=0.001). The mean normalized synergy increased as the number of recurrent edges increased (0.0080 [0.0069 0.0093] vs. 0.012 [0.010 0.014] vs. 0.014 [0.012 0.017] for 0, 1, and 2 edges, respectively; Fig 4B), reflected by a significant positive correlation between normalized synergy and number of recurrent edges (Spearman *r* = 0.25, n=675, p<1×10^−8^). The main effect of feedback was also significant (*F*(2,148)=50, p<1×10^−17^). The mean normalized synergy decreased as the number of feedback edges increased (0.012 [0.011 0.014] vs. 0.011 [0.009 0.013] vs. 0.008 [0.007 0.010] for 0, 1, and 2 edges, respectively; Fig 4C), reflected by a significant negative correlation between normalized synergy and number of feedback edges (Spearman *r* = −0.22, n=675, p<1×10^−6^). There was also a significant interaction between recurrence and feedback on normalized synergy (*F*(4,296)=6.7, p<1×10^−4^). Specifically, normalized synergy was highest in motifs with the most recurrent edges and the fewest feedback edges, and normalized synergy was lowest in motifs with the fewest recurrent edges and the most feedback edges. Taken together, these results show that, across these networks, motifs with greater upstream recurrence have greater normalized synergy, and motifs with greater feedback have lesser normalized synergy.

**Fig 4.**
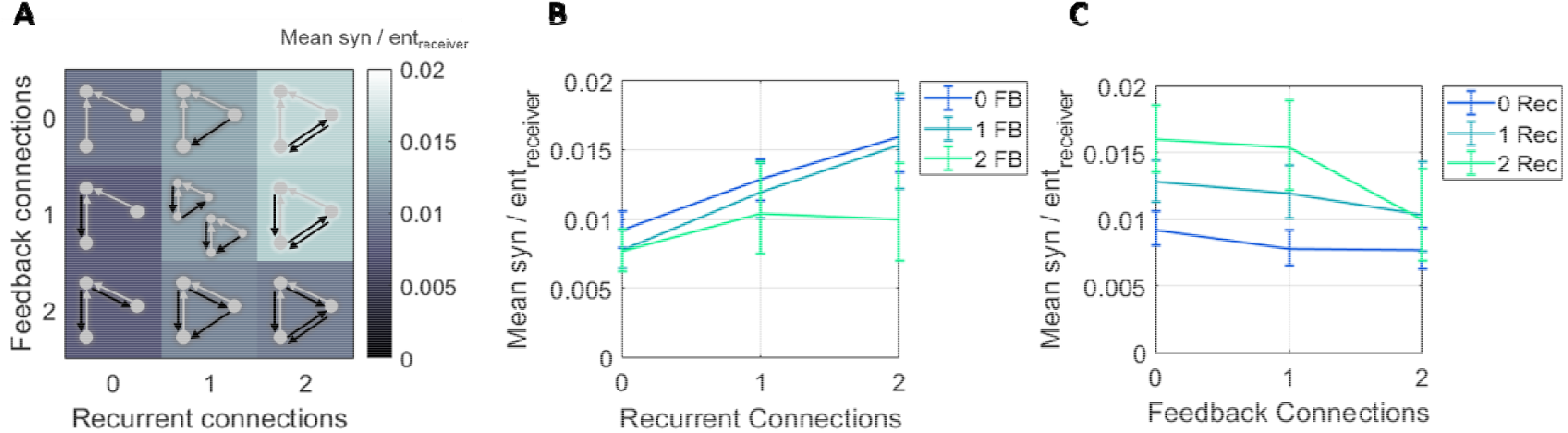
Normalized synergy increased with greater recurrence and decreased with greater feedback. (A) Motifs are ordered based on the number of recurrent edges (columns) and feedback edges (rows). The background heatmap, wherein brighter colors reflect larger normalized synergy values, replots the central tendency of the values shown in Figure 3. (B) Curves representing rows shown in A, plotted with errorbars computed across networks, show that synergy increased as the number of recurrent edges increased. (C) Curves representing columns shown in A, plotted with errorbars computed across networks, show that synergy decreased as the number of feedback edges increased. Errorbars are 95% bootstrap confidence intervals around the mean.

To understand if/how the normalization of synergy by the receiver entropy affected these results, we separately examined how raw synergy (i.e., non-normalized synergy) varied across motifs (Fig 5). As with normalized synergy, raw synergy was positively correlated with the number of recurrent edges (Spearman *r* = 0.15, n=675, p<0.001). A two-way ANOVA examining how raw synergy values varied across levels of recurrence and feedback found the main effect of recurrence trending toward significance (*F*(2,148)=2.6, p=0.08; Fig5B). The results for feedback, however, were qualitatively different when examining raw synergy versus normalized synergy. Raw synergy also increased with the number of feedback edges (Spearman *r* = 0.10, n=675, p=0.01) rather than the decrease that was observed for normalized synergy. The main effect of feedback in the two-way ANOVA was significant (*F*(2,148)=4.7, p=0.01; Fig 5C). We again found a significant interaction between recurrence and feedback levels (*F*(4,296)=7.8, p<1×10^−5^) such that the contribution of recurrence and feedback to raw synergy failed to sum linearly. Thus, consistent with our previous findings [6] these results indicate that raw synergy generally increases with greater edge density.

**Fig 5.**
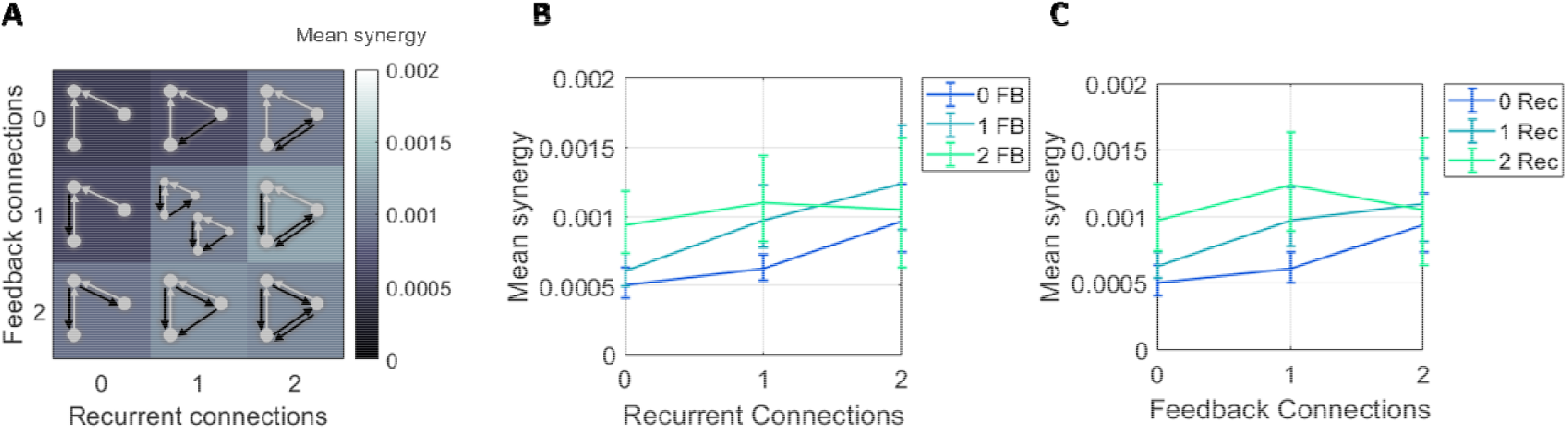
Raw synergy increased with greater recurrence and greater feedback. (A) Mean synergy increased with the number of recurrent and feedback edges in motifs. (B) Curves representing rows shown in A, plotted with errorbars computed across networks, show that synergy increased as the number of recurrent edges increased, although the difference in means across levels was not significant. (C) Curves representing columns shown in A, plotted with errorbars computed across networks, show that synergy increased as the number of feedback edges increased. Errorbars are 95% bootstrap confidence intervals around the mean.

The fact that raw synergy and normalized synergy varied in opposite directions with respect to the number of feedback edges reveals that the normalizing term, receiver entropy, mediated the relationship between feedback and synergy. Receiver entropy differed significantly across motifs (Fig. 6). A two-way ANOVA found significant main effects of recurrence (*F*(2,148)=32.7, p<1×10^−11^) and feedback (*F*(2,148)=15.1, p<1×10^−5^) and a significant interaction effect (*F*(4,296)= 13.19 p<1×10^−9^). Receiver entropy was negatively correlated with recurrence level (Spearman *r* = −0.16, n=675, p<0.001; Fig 6B) and strongly positively correlated with feedback level (Spearman *r* = 0.41, n=675, p<1×10^−22^; Fig 6C). The finding that there was greater receiver entropy in motifs with more feedback edges accounts for the different pattern of results observed for raw and normalized synergy across feedback levels. Motifs with greater feedback, these results suggest, emerge around receiver nodes with greater entropy. While these motifs generate more raw synergy, that synergy accounts for a lesser proportion of the entropy of the respective receiver (i.e., less normalized synergy). Recurrence, however, did not display such a dependence. Whether normalized by receiver entropy or not, synergy was positively related to the number of recurrent edges in the motif.

**Fig 6.**
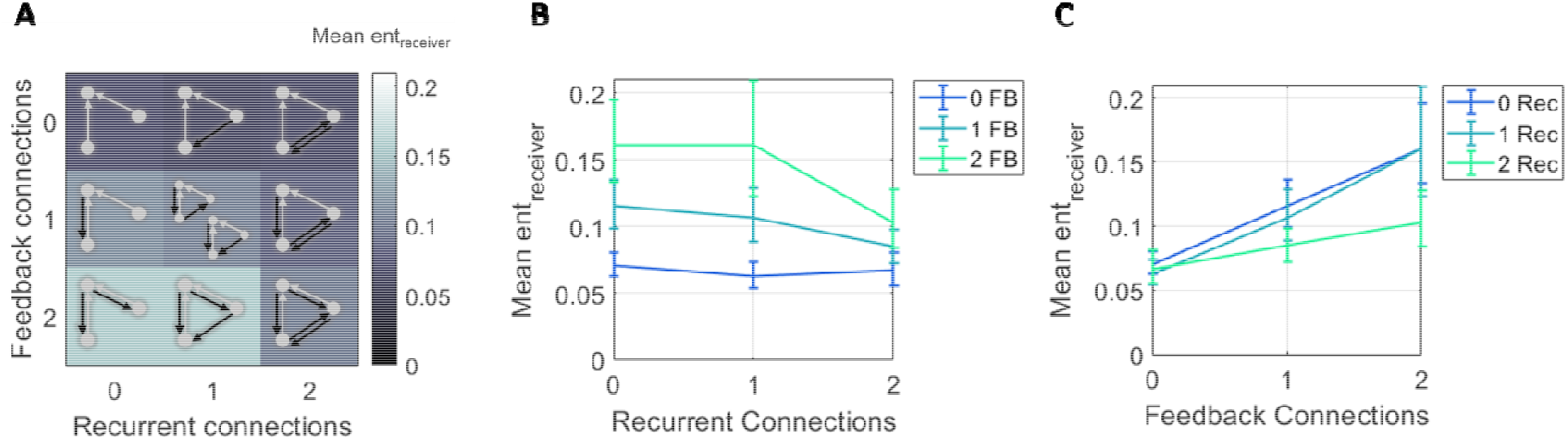
Receiver entropy decreased with greater recurrence and increased with greater feedback. (A) Mean entropy decreased with the number of recurrent edges and increased with the number of feedback edges in motifs. (B) Curves representing rows shown in A, plotted with errorbars computed across networks, show that entropy decreased as the number of recurrent edges increased. (C) Curves representing columns shown in A, plotted with errorbars computed across networks, show that entropy increased as the number of feedback edges increased. Errorbars are 95% bootstrap confidence intervals around the mean.

Next, we sought to determine whether the observed relationship between normalized synergy and motif structure could be the product of systematic variance across motifs in either the strength of feedforward edge weight or the mutual information between the senders. Both are predictive of normalized synergy [6,10]. To address each potential confound, we regressed out the variance associated with each factor and asked if the residual normalized synergy was nonetheless correlated to the motif structure (Table 1). The residual normalized synergy remained significantly positively related to the number of recurrent edges whether we regressed out variance associated with feedforward edge weight (*F*(2,148)=9.4, p<1×10^−3^; Spearman *r* = 0.37, n=675, p<1×10^−18^) or the mutual information between the senders (*F*(2,148)=29.4, p<1×10^−10^; Spearman *r* = 0.49, n=675, p<1×10^−33^). Thus, the positive relationship between normalized synergy and motif recurrent edge number is not accounted for by covariance in either feedforward edge weight or mutual information between senders.

**Table 1.** Relationships between residual normalized synergy and recurrence and feedback, after regressing out potential confounding covariates: strength of feedforward edge weights (Syn-FF), and mutual information between senders (Syn-MI). Columns 1-3 (*df, F, p*_ANOVA_) show the results of a repeated measures ANOVA for the residual normalized synergy predicted by the number of recurrent and feedback edges. Columns 4-5 (*rho, p_rho_*) show the results of Spearman rank correlations between residual normalized synergy and the number of recurrent and feedback edges. P-values significant at the α=0.05 level are in bolded font.

| **Syn-FF** | *df* | *F* | *p*<sub>ANOVA</sub> | *rho* | *p*<sub>rho</sub> |
| --- | --- | --- | --- | --- | --- |
| Recurrent | 2, 148 | 9.4 | **<1x10<sup>-3</sup>** | 0.37 | **<1x10<sup>-18</sup>** |
| Feedback | 2, 148 | 1.0 | 0.36 | -0.11 | **0.01** |
| Recurrent x Feedback | 4, 296 | 3.4 | **0.01** | -- | -- |
| **Syn-MI** | *df* | *F* | *p*<sub>ANOVA</sub> | *rho* | *p*<sub>rho</sub> |
| Recurrent | 2, 148 | 29.4 | **<1x10<sup>-10</sup>** | 0.49 | **<1x10<sup>-33</sup>** |
| Feedback | 2, 148 | 14.5 | **<1x10<sup>-5</sup>** | -0.28 | **<1x10<sup>-10</sup>** |
| Recurrent x Feedback | 4, 296 | 2.8 | **0.03** | -- | -- |

The negative relationship between normalized synergy and motif feedback edge number, likewise, was not affected by regressing out variance associated with mutual information between senders (*F*(2,148)=14.5, p<1×10^−5^; Spearman *r* = −0.28, n=675, p<1×10^−10^). The residual normalized synergy, after regressing out variance associated with feedforward edge weight, maintained a significant negative correlation with motif feedback edge count (Spearman *r* = - 0.11, n=675, p=0.01) but was no longer a significant main effect in the two-way ANOVA (*F*(2,148)=1.0, p=0.36). This loss of significance also occurred without the normalization by receiver entropy (*F*(2,148)=0.7, p=0.50). The sensitivity of the relationship between synergy and motif feedback edge count to regressing out variance associated with feedforward edge weight indicates variance in synergy associated with the number of feedback edges could be accounted for by variance in feedforward edge strength. Indeed, in a post hoc test, we found that feedforward edge weight was significantly negatively correlated with the number of feedback edges (Spearman *r* = −0.22, n=675, p<1×10^−6^). As described further in the discussion, the negative correlation between feedforward and feedback information flow leaves ambiguity to be resolved by future work regarding the separate effects of feedback and feedforward information flows on synergistic integration. What can be said is that synergy was greater when the ratio of feedforward to feedback information flow was larger (Spearman *r* = 0.50, n=675, p<1×10^−35^).

Extending our analysis of how normalized synergy is related to the *number* of recurrent and feedback edges to ask if it is similarly related to the *strength* of these edges, we performed a multiple linear regression for each network, using the strength of feedforward, recurrent, and feedback edge weights as predictors of normalized synergy. We then analyzed the distribution of beta coefficients for each term (Fig 7). As expected from prior work [6], feedforward edge weight was a significant predictor of normalized synergy (z_s.r_. = 7.52, n=75, p<1×10^−13^). Recurrent edge weight was also a significant predictor of normalized synergy (z_s.r_. = 6.12, n=75, p<1×10^−9^). Conversely, feedback edge weight was not a significant predictor of normalized synergy (z_s.r_. = −0.93, n=75, p=0.35). To determine if motif structure could account for variance beyond what could be accounted for by the edge weights, we performed an ANOVA on the residuals across networks. We observed a significant main effect of the number of recurrent edges (*F*(2,148)=3.2, p=0.04), which remained positively correlated with the residual normalized synergy (Spearman *r* =0.17, n=675, p<1×10^−4^). We observed no significant main effect of the number of feedback edges (*F*(2,148)=0.8, p=0.93) but did find a marginally significant negative correlation between feedback edge number and residual normalized synergy (Spearman *r* =-0.08, n=675, p=0.05). We observed no significant interaction effect (*F*(4,296)=1.96, p=0.10). These results show that recurrence is positively related to synergistic integration when quantified as the weight of the motif edges as well as the number of edges. The same could not be said for feedback edge weight which was not significantly correlated to synergistic integration.

**Fig 7.**
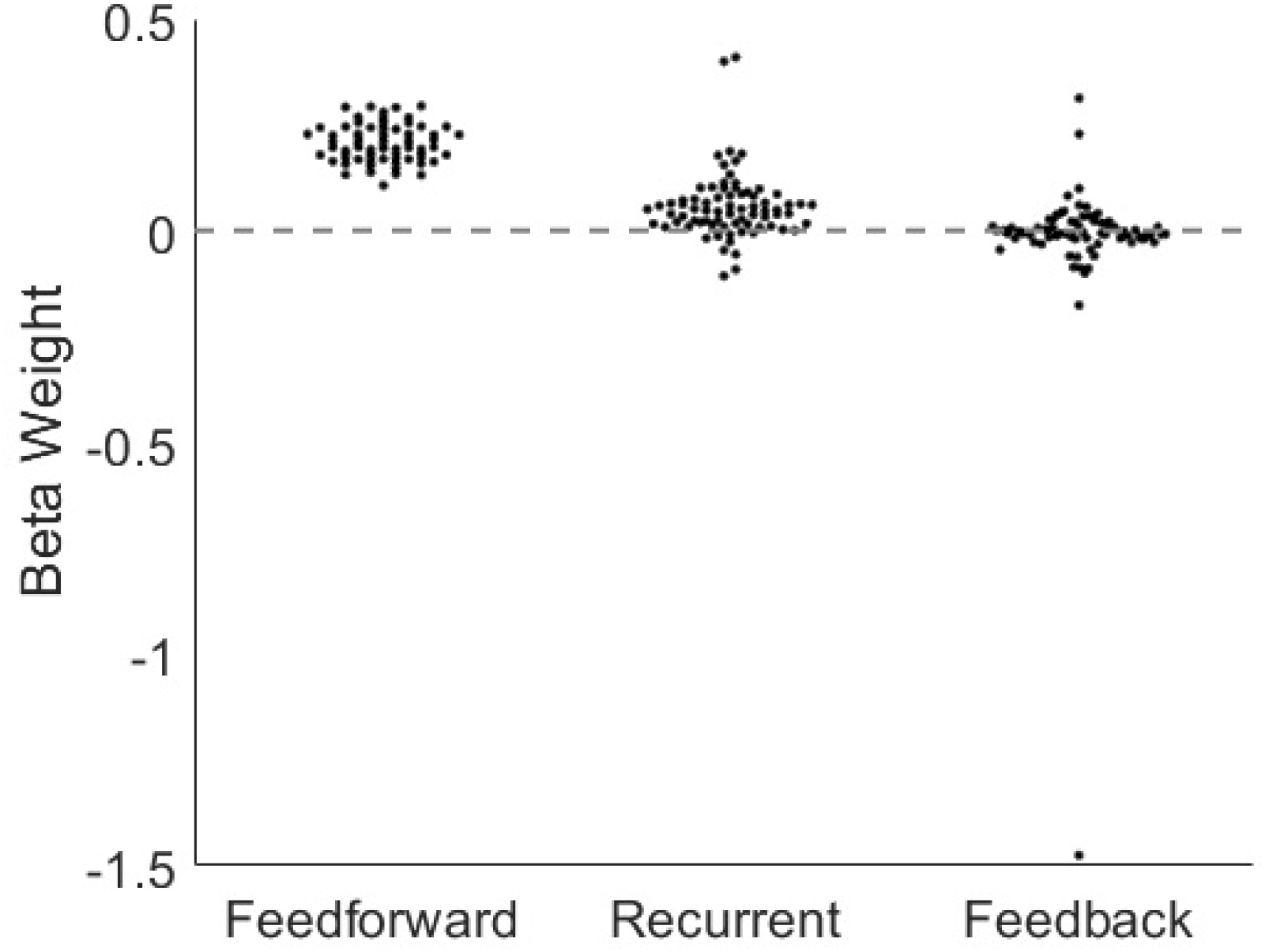
The strength of feedforward and recurrent edges was predictive of normalized synergy. Histograms of beta weights for multiple linear regressions performed on the networklevel revealed that feedforward and recurrent edge weights were reliable predictors of normalized synergy, across networks. Feedback edge weight was not a reliable predictor of normalized synergy.

The above results show that the number of recurrent edges among upstream neurons contributed a novel source of synergy beyond the variance accounted for by the receiver entropy and by the weight of the feedforward edges. This result was not sensitive to the type of normalization performed (S6 Fig and S7 Fig), nor was it sensitive to the exact implementation of PID used (S13 Fig and S14 Fig). The observed effects on synergy were consistent with known relationships between synergy, redundancy, and multivariate TE at these timescales (S8-S12 Fig). The observed effects also held when the three timescales are analyzed separately (S15 Fig).

### Recurrent and feedback motifs are rare but overrepresented

To gain perspective as to how our findings regarding the influence of recurrence and feedback on synergy relate to network-wide processing, we asked how prevalent each type of effective connectivity was in our networks. To do this, we calculated the percentage of networkwide triads accounted for by each motif (Fig 8).

**Fig 8.**
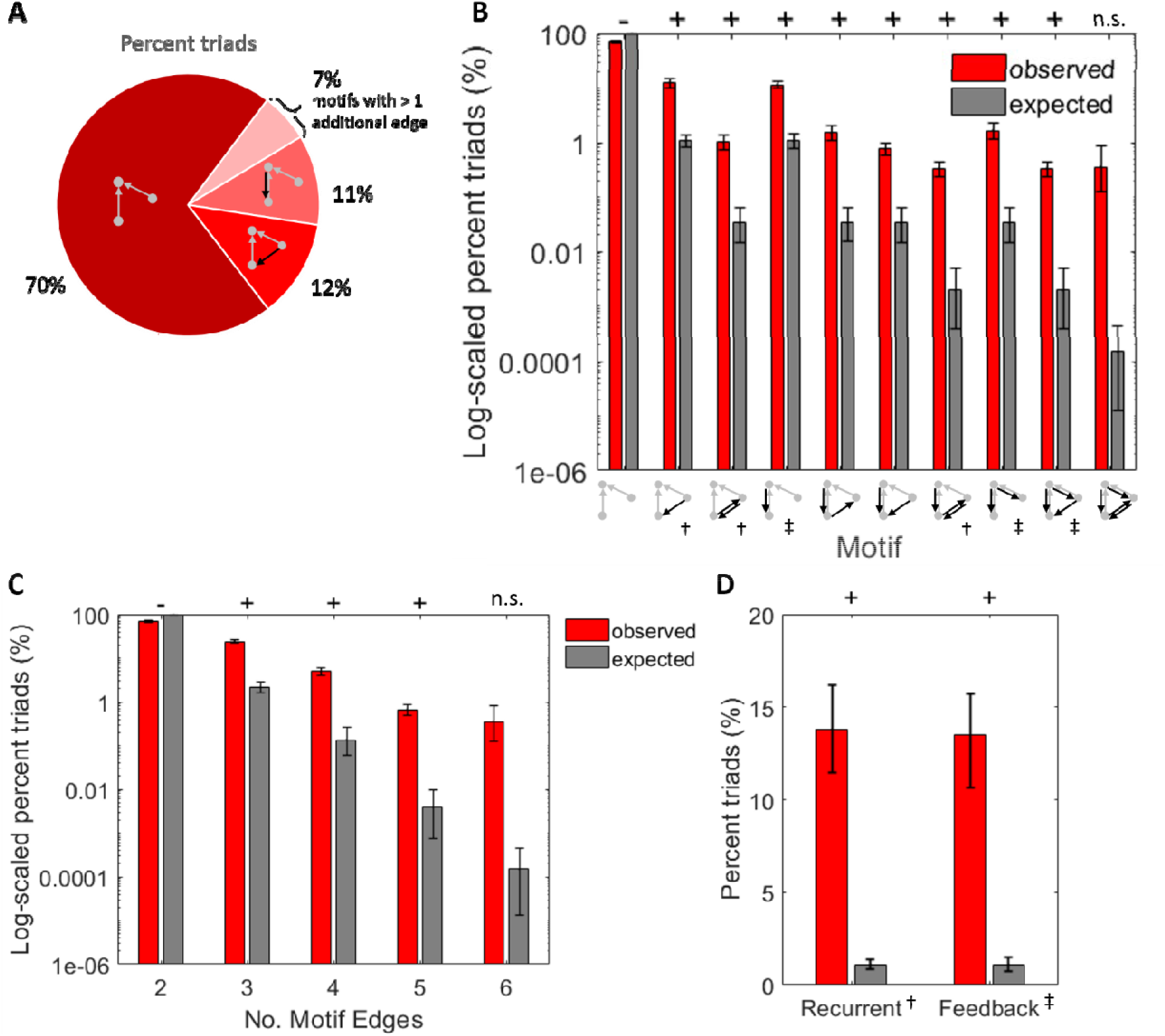
Recurrent and feedback motifs were rare but occurred more than expected given the network-wide edge density. (A) Percent of network triads accounted for by each motif type. Motifs with greater edge density were more rare. Values indicate means across all networks. (B) Log10 scaled observed percentages of triads compared to expected percentages of triads per motif. Expected percentages obtained by raising the probability of observing a connection to the power of the number of edges in the motif, for each network. (C) Log_10_ scaled observed and expected percentages of triads in (B), grouped by the number of edges they contain. (D) Linearly scaled observed and expected percentages of triads in (A), grouped by the number of recurrent and feedback edges they contain. Significance indicators: ‘+’ indicates significantly more than expected and ‘-’ indicates significantly less than expected. For all significant values, *p<1×10^−6^*. Motif indicator: † Recurrent, ‡ Feedback.

Consistent with the sparsity of these networks (average edge density: 1.14% [0.83% 1.54%]), the incidence of each motif decreased rapidly as a function of the number of edges contained in the motif. The first (i.e., default) motif, containing only 2 feedforward edges, was most prevalent and accounted for 70.12% [67.15% 73.07%] of the observed synergistic 3-node triads (Fig 8A-C). Motifs with 3 edges, whether recurrent or feedback, accounted for 23.94% [21.76% 26.14%] of the synergistic 3-node triads (Fig 8B-C). This is significantly greater than the 2.12% [1.60% 2.83%] that would be expected by chance given random networks with the same sparsity (t = 21.24, n = 75, p < 1×10^−32^). Motifs with 4, 5, and 6 edges were similarly overrepresented from what would have been expected in random networks, but progressively decreased in prevalence (4 edge motifs: 4.93% [4.03% 5.92%] vs. 0.13% [0.06% 0.26%], t = 10.27, n = 75, p < 1×10^−15^; 5 edge motifs: 0.66% [0.47% 0.88%] vs. 0.0039% [0.0008% 0.01%], t = 6.39, n = 75, p < 1×10^−7^; 6 edge motifs: 0.36% [0.12% 0.87%] vs. 0.0002% [0.0000% 0.0004%], t = 1.99, n = 75, p = 0.051; Fig 8C). These results agree with findings in similar networks generated from the same data [46]. Finally, recurrent and feedback motifs occurred significantly more than would have been expected in random networks (recurrent: 13.79% [11.52% 16.28%] vs. 1.10% [0.82% 1.48%], t = 10.68, n = 75, p < 1×10^−15^; feedback: 13.47% [11.41% 15.70%] vs.1.10% [0.82% 1.48%], t = 10.97, n = 75, p < 1×10^−16^; Fig 8D). Importantly, all motifs with recurrent and feedback edges, with the exception of the 6-edge motif, occurred more frequently than expected given network connection densities. Thus, the sparsity of our networks did not preclude our ability to detect recurrent and feedback motifs.

To test for evidence of selection bias toward or away from motifs with recurrent or feedback edges, we tested whether one type was more or less prevalent among the motifs containing a given number of edges. The null distribution would be an equal number of each. Among 3-edge synergistic motifs, the extra edge was recurrent in 50.7% [44.2% 57.1%] of the triads. This was not significantly different from 50% (t = 0.22, n = 75, p = 0.83). Likewise, across triads with 4 and 5 edges, we found no evidence of bias toward one type of effective connectivity versus the other. In 4-edge motifs with both additional edges being the same, both additional edges were recurrent in 43.1% [35.1% 51.1%] of triads (t = −1.67, n = 75, p = 0.10) and both additional edges were feedback in 56.9% [48.7% 64.7%] of triads (t = 1.67, n = 75, p = 0.10). In 4 edge motifs with both additional edges being different, 1 additional edge was recurrent and 1 was feedback in 49.1% [43.3% 54.5%] of triads (t = −0.29, n = 75, p =0.76). In 5-edge motifs, the additional edge was recurrent in 50% [50% 50%], t = 0, n = 75, p = 1; 6-edge motifs were not included in this as they contain the same number of feedback and recurrent edges by definition). Consistent with this, there was also no difference in the overall likelihood of observing a recurrent versus a feedback edge (7.5% [6.4% 8.7%] vs. 7.6% [6.6%, 8.8%]; z_s.r_.= −0.07, n=75, p=0.94).

Given the similar incidence of motifs containing recurrent and feedback edges, but significant differences in the synergy observed for each motif type, synergistic triads containing recurrent edges could be expected to account for a larger percentage of the network-wide synergy (Fig 9). Network-wide synergy was the sum of synergy, taken across all triads in a network. Indeed, recurrent motifs comprised 13.79% [11.52% 16.28%] of triads and accounted for 20.43% [17.26% 23.85%] of network-wide synergy. Feedback motifs comprised 13.47% [11.41% 15.70%] of triads and only 10.12% [8.37% 12.11%] of network-wide synergy (Fig 9A, inset). Thus, although recurrent and feedback motifs accounted for similar percentages of network triads (z_s.r_= 0.09, n=75, p=0.92), recurrent motifs accounted for a significantly higher percentage of network synergy than feedback motifs (z_s.r_= 4.18, n=75, p<1×10^−4^).

**Fig 9.**
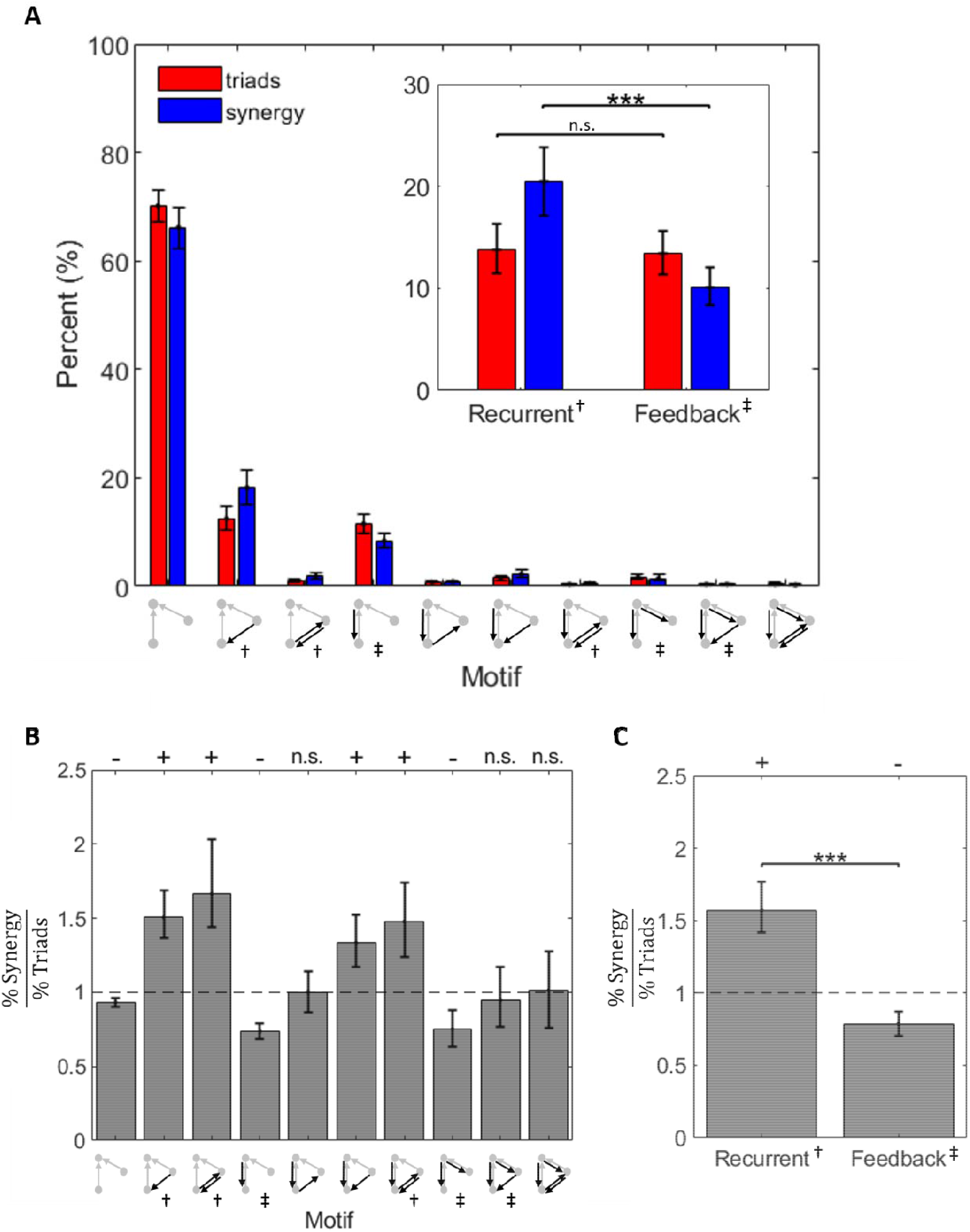
Recurrent and feedback motifs accounted for more and less network-wide synergy than expected, respectively. (A) Both recurrent motifs and feedback motifs were relatively rare, and they accounted for a relatively small proportion of overall synergy. Red bars from Fig 8B are replotted on a linear scale here for comparison. Inset: Recurrent motifs were as common as feedback motifs, but they accounted for significantly more synergy than feedback motifs. Red bars from Fig 8D are replotted here for comparison. (B) The ratio of percent synergy to percent triads is shown per motif. Values above one indicate that the motif accounts for greater network-wide synergy than it does triads. Values less than one indicate that the motif accounts for less network-wide synergy than it does triads. (C) Recurrent motifs accounted for more synergy than expected given their frequency. Conversely, feedback motifs accounted for less synergy than expected given their frequency. Significance was determined by asking whether the distribution of ratios for each motif came from a distribution whose mean is equal to 1 (t-test). Significance indicators: ‘+’ indicates significantly more than expected and ‘-’ indicates significantly less than expected. For all significant values,*p<0.001*. Central tendency shown in each figure is mean and error bars are 95% bootstrap confidence intervals around the mean. Mean was selected over median to ensure that percentages sum to 100. Significance indicator: *** p<0.001. Motif indicator: † Recurrent, ‡ Feedback.

To determine whether motifs accounted for more synergy than expected given their frequency, we calculated the ratio of percent synergy to percent triads for each motif (Fig 9B-C). Values greater than one indicated that the motif accounted for more synergy than expected given its frequency. Values less than one indicated that the motif accounted for less synergy than expected given its frequency. We observed that recurrent motifs accounted for significantly greater network-wide synergy than expected given their frequency (z_s.r_. = 6.28, n=75, p<1×10^−9^), and feedback motifs accounted for significantly less network-wide synergy than expected given their frequency (zs.r. = −4.35, n=75, p<1×10^−4^; Fig 9C).

## Discussion

Understanding the relationships between directed information flow (feedforward, feedback, and upstream recurrence) and synergistic processing in cortical networks is essential for understanding how neural networks compute. We previously showed that synergistic processing varies directly with feedforward information flow [6]. Here, we examined the influence of recurrent and feedback information flow on synergistic information processing in organotypic cortical cultures. Using information theoretic and network analyses of the spiking activity of hundreds of simultaneously recorded neurons from organotypic cultures of mouse somatosensory cortex, we showed for the first time that recurrent and feedback information flow in functional local microcircuits varies systematically with the amount of synergy observed in those microcircuits. Specifically, we found that greater recurrence in motifs predicted greater synergy while greater feedback predicted lesser synergy (Figure 10). Interestingly, the strength of feedforward effective connections, a covariate of synergy [6], could account for much of the variance associated with the feedback-synergy relationship. It could not, however, account for the recurrence-synergy relationship (Table 1). Thus, recurrence predicted synergistic processing above and beyond that predicted by the strength of inputs. Additionally, we found that, although recurrent motifs were somewhat rare in our networks--comprising 14% of all synergistic motifs-- they account for 20% of the total network-wide synergy. Feedback motifs--comprising 13% of all synergistic motifs--were roughly matched for prevalence with recurrent motifs, but only accounted for 10% of the total network-wide synergy. Thus, with similar prevalence, recurrent motifs accounted for twice as much synergy as feedback motifs.

**Fig 10.**
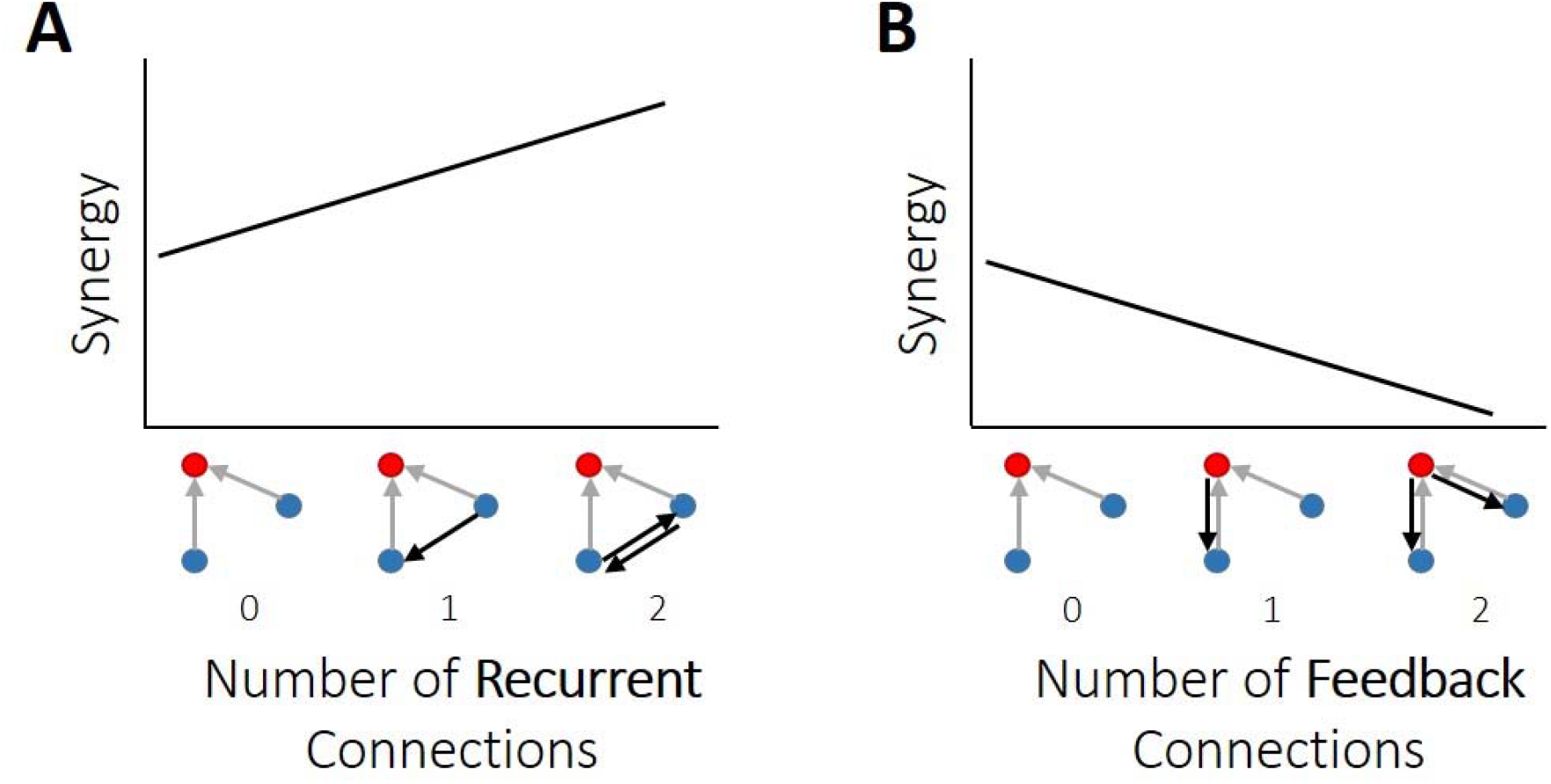
Summary of findings regarding how synergy is related to recurrent and feedback information flow in organotypic cultures of mouse cortex. (A-B) Synergy had a positive relationship with the number of recurrent edges and a negative relationship with the number of feedback edges. That is, synergy was elevated where there was greater upstream recurrent information flow. Synergy was diminished where there was greater feedback information flow.

### Relationship to previous work

Our finding that synergy increased with greater recurrence is consistent with previous work showing that recurrent connections—both functional and structural—are necessary for pattern completion tasks, both in biological [12,13,15-17,19,20] and artificial networks [14,19]. Such tasks involve the integration of multiple, distinct features to generate a coherent representation, a process that involves some form of synergistic processing. Our finding that synergy decreased with greater feedback agrees with theoretical frameworks [33–36] and experimental studies [29-31,37,47] suggesting that feedback processes serve to reduce the variance with which lower-level neurons can account for variance in higher-level neurons, thereby reducing the strength of feedforward information flow, and resulting in reduced synergy.

Our finding that increased recurrent connectivity corresponded to greater synergistic processing is also consistent with previous analyses of the topological determinants of synergistic processing in functional networks of cortical cultures. For example, one such analysis found that synergistic processing was directly related to the ‘out-degree’ of effective connectivity of the upstream neurons [8]. That is, the more neurons that a given upstream neuron made effective connections with, the greater the resulting synergy was in the recipient neurons. Similarly, we have previously shown that neurons in functional rich clubs of cortical microcircuits (i.e., highly-intercommunicating neurons) do about twice as much synergistic processing as neurons outside of the rich clubs [6]. We have also shown previously that greater similarity (i.e. synchrony) of transmitters, such as might be generated by strong intercommunication, predicts greater synergy at synaptic timescales [10].

The strength of feedforward information flow was an important consideration here when analyzing the relationship between the number of recurrent/feedback edges in a motif and synergy. We have shown previously that the strength of feedforward information flow is a strong, positive predictor of the amount of synergy [6]. Here, we performed a control analysis where this relationship was first regressed out of the synergy values before asking whether recurrence and/or feedback were predictive of synergy. The positive relationship between recurrence and synergy persisted after regressing out the influence of feedforward information flow, indicating that recurrent information flow reflects a novel source of explanatory power over synergy. We hypothesize that this additional synergy emerges because recurrent information flow increases the capacity of the transmitter neurons to jointly predict the behavior of the receiver, resulting in more synergy than if it just increased the amount of bivariate transfer entropy. The relationship between the number of feedback edges and synergy, however, was lost after regressing out variance associated with feedforward information flow. This suggests that feedforward and feedback information flow account for common variance in the observed synergy as discussed in greater detail in the next section.

### Variability of the synergy-feedback relationship

While the relationship between synergy and motif recurrent edge count was robust across all analyses we performed, the relationship between synergy and motif feedback edge count varied across analyses, exhibiting both negative and positive relationships with synergy. First, we found that it mattered whether raw synergy (i.e., total synergy without scaling of any sort) or normalized synergy (i.e., raw synergy normalized by the entropy of the receiving neuron so as to indicate the percentage of the entropy accounted for) was examined. Raw synergy was positively correlated with motif feedback edge count while normalized synergy was negatively correlated with motif feedback edge count. We found that this reversal could be accounted for by the fact that the receiver entropy, the normalizing term, itself varied significantly as a function of motif feedback edge count. The receivers in motifs with more feedback edges had more entropy. Thus, while greater raw synergy emerges in these motifs, it accounts for a lower percentage of the receiver entropy.

Second, we found that regressing out variance in synergy, whether normalized or not, associated with feedforward information flow resulted in a loss of significance of the relationship to motif feedback edge count. As noted above, this result indicates that variance in synergy associated with the number of feedback edges was also associated with variance in feedforward information flow. Two scenarios could yield this result. In both, feedforward and feedback information flows have opposing effects on synergy. These scenarios differ with regard to whether feedforward and feedback flows are independent. If independent, then feedforward and feedback processing do not directly affect each other, but simply have contrasting effects on synergy. Our finding of a significant negative correlation between these terms, however, argues against such a scenario. Rather, this negative correlation suggests that feedforward information flow affects the amount of feedback information flow and/or vice versa.

### Influence of network edge density

Network edge density was another important consideration in studying the influence of recurrent or feedback edges on synergy. Our networks were sparse, consistent with densities observed in previous studies of biological neural networks [43,48-51]. With this sparsity, it is possible that our results might have been skewed by the lack of connectivity, which would translate to a lack of observations for motifs with greater connectivity (i.e. recurrent and feedback motifs). We investigated the influence of sparsity on our results by asking how the expected frequency of motifs, given the probability of a single connection, compared to the frequency of motifs that we observed in our networks. We found that our networks had significantly more instances of both recurrent and feedback motifs than expected by chance given the baseline probability of observing a significant edge. Thus, we concluded that the sparsity of our networks did not curtail our ability to observe these motifs. Moreover, the fact that recurrent and feedback motifs occurred more than expected by chance may indicate that such motifs, which evolve from network dynamics, are important for network processing.

### Use of organotypic cultures

This work required the ability to record the spiking activity of hundreds of neurons simultaneously. This was made possible by our use of organotypic cultures. While organotypic cultures naturally differ from intact in vivo tissue, organotypic cultures nonetheless exhibit synaptic structure and electrophysiological activity very similar to that found in vivo [52–58]. For example, the distribution of firing rates observed in cultures is lognormal, as seen in vivo [59], and the strengths of functional connections are lognormally distributed, similar to the distribution of synaptic strengths observed in patch clamp recordings [reviewed in 60,61]. However, neural cultures exhibit bursty dynamics that evolve as a function of the age of the culture [62–65]. Here, recordings were collected between 2 and 4 weeks after culture preparation, corresponding to an age at which ‘broad network bursts’ (2s) are commonly observed [63]. Evidence regarding the existence of multi-second ‘network bursts’ in the intact brain, outside of inactive conditions [66], is limited. Recordings from intact brains *in situ* do show, however, that brief local bursts occur regularly and are considered behaviorally relevant [67–70]. Observed differences between cultures and the intact brain regarding the spatiotemporal properties of bursts likely reflect differences in physiological condition [71–73]. Yet, even with these differences, the most parsimonious hypothesis is that the relationships observed between functional dynamics and informational metrics (e.g., synergy) observed in neural cultures, as done here, are informative for understanding brain function. Nonetheless, additional work will need to be done to understand how the relationships between synergy and recurrence and synergy and feedback observed in vitro differ from what may exist in vivo, particularly in the context of behavior.

### Co-occuring spiking dynamics

The results obtained here were the product of the recorded spiking dynamics. The full nature and scope of this dependence is a topic of active investigation [6,8,10,59,67]. To facilitate recognition of factors of key relevance as new work with convergent or divergent results emerge, we review briefly here the spiking dynamic properties, as characterized previously, for this dataset. Neurons in our recordings had a mean firing rate of 2.1 Hz [2.0 Hz, 2.2 Hz] and generated rhythmic bursts of activity [41], with cross-correlation peaks in the frequency bands corresponding to theta (4-12 Hz), beta (12-30 Hz), gamma (30-80 Hz), and high frequency (100-1000 Hz) oscillations [42]. Bursts typically lasted between 1 and 10 seconds, and were typically separated by 10 seconds, though significantly shorter durations and intervals occurred [41]. Despite exhibiting bursts, neuron spiking was relatively sparse, with 80% of neurons firing at <3 Hz. These spiking dynamics are well aligned with those observed *in vivo* [67,74].

The sparsity of neuron firing was likely an important factor in combination with the methods used here. The low neuron firing rates together with the construction of our timescales, which binned neuron spiking into 1 ms, 1.6 ms, and 3.5 ms bins–1000 bins/sec, 625 bins/sec, and 286 bins/sec, respectively–resulted in spike trains in which a large majority of bins contained 0 spikes. An impact of this sparsity is that the TE calculations had greater sensitivity for detecting excitatory interactions. Therefore, we cannot speak to the influence of inhibitory activity in producing the results shown here. Future work should aim to determine the relative influence of excitation and inhibition on synergistic integration.

### Relevance of spontaneous activity

While stimulus-driven activity has been favored in research for its ability to provide insight into neural coding mechanisms, such studies assume that the brain is primarily reflexive and that internal dynamics are not informative with regard to information processing. However, internally-driven spontaneous activity of neurons, or activity that does not track external variables in observable ways, has been repeatedly shown to be no less cognitively interesting than stimulus-linked activity [75,76; for a review see 77,78]. Not only is spontaneous activity predominant throughout the brain, but it also drives critical processes such as neuronal development [11,79,80].

### Limitations of our PID approach

Multiple methods exist for calculating synergy [9,81-83] and, here, we used PID as described by [7]. We chose this measure because it is capable of detecting linear and nonlinear interactions and it has been shown to be effective for our datatype [6,8,10]. In addition, unlike other methods [84,85], PID of mvTE can decompose the interaction into non-negative and nonoverlapping terms. However, there is the reasonable concern that PID overestimates the redundancy term and consequently synergy [81–83]. Here, to address this issue, we also implemented an alternate form of PID that estimates synergy based on the lower bound of redundancy. The result of this, in the present context, is that the effective threshold for triads to generate synergy is higher. Nonetheless, we found that this approach yielded the same qualitative pattern of results (S13 Fig, S14 Fig). Because the synergy-recurrence relationship holds when assessed using both the upper and lower bounds of synergy, we believe the relationship will hold for alternative implementations of PID that determine intermediate levels of synergy.

The present work did not examine interactions larger than triads due to the multi-fold increase in the computational burden that arises in considering higher order synergy terms. In addition to the combinatorial explosion of increased numbers of inputs, the number of PID terms increases rapidly as the number of variables increases. However, based on bounds calculated for the highest order synergy term by [8], it was determined that the information gained by including an additional input beyond two either remained constant or decreased. From this, they inferred that lower order (two-input) operations dominated. Nonetheless, further investigation of this point will be worthwhile as improvements in computational wherewithal enable it.

### Significance of the present work

This work could inform future research on the importance of the topology of biological networks. The functional topology of biological neural networks has already been shown to influence neural information processing [6,8,59]. The present results add to our growing understanding of how the structure of neuronal interactions shapes neuronal behavior. These findings could also inform further research on artificial intelligence. Specifically, our results could be used in applied efforts to design engineered systems for the optimization of computational power and efficiency.

### Conclusion

In summary, the present study demonstrated that, in functional networks observed in organotypic cultures, recurrent information flow between senders in 3-neuron synergistic motifs is positively related to the synergistic integration by the receiving neuron. We also showed that feedback information flow from the receiving neuron to the senders in 3-neuron synergistic motifs is negatively related to the amount of synergistic integration by the receiving neuron. Although the frequencies of motifs with relatively more recurrent or feedback information flow are matched, they account for more and less synergy than expected, respectively. These results add to a growing body of work regarding the interdependence of synergistic integration and functional network topology. Taken together, these findings provide increasing evidence of the influence of recurrent and feedback information flow on overall neural network information processing.

## Materials & Methods

To answer the question of how computation is related to feedback and recurrence in cortical circuits, we combined network analysis with information theoretic tools to analyze the spiking activity of hundreds of neurons recorded from organotypic cultures of mouse somatosensory cortex. Here we provide an overview of our methods and focus on those steps that are most relevant for interpreting our results. A comprehensive description of all our methods can be found in the Supporting Information.

All procedures were performed in strict accordance with guidelines from the National Institutes of Health, and approved by the Animal Care and Use Committees of Indiana University and the University of California, Santa Cruz.

### Electrophysiological recordings

All results reported here were derived from the analysis of electrophysiological recordings of spontaneous activity from 25 organotypic cultures prepared from slices of mouse somatosensory cortex between 2 and 4 weeks after culture preparation. One hour long recordings were performed at 20 kHz sampling using a 512-channel array of 5 μm diameter electrodes arranged in a triangular lattice with an inter-electrode distance of 60 μm (spanning approximately 0.9 mm by 1.9 mm). Once the data were collected, spikes were sorted using a PCA approach [41-42,86] to form spike trains of between 98 and 594 (median = 310) well isolated individual neurons depending on the recording.

### Network construction

Networks of effective connectivity, representing global activity in recordings, were constructed following the methods described by [8,41]. Briefly, weighted effective connections between pairs of neurons were established using transfer entropy (TE)[87]. To consider synaptic interactions, we computed TE at three timescales spanning 0.05 - 14 ms, discretized into overlapping bins of 0.05-3 ms, 1.6-6.4 ms, and 3.5-14 ms, resulting in 75 different networks. Only significant TE, determined through comparison to the TE values obtained with jittered spike trains (α = 0.001; 5000 jitters), were used in the construction of the networks. TE values were normalized by the total entropy of the receiving neuron so as to reflect the proportion of the receiver neuron’s capacity that can be accounted for by the transmitting neuron. Note, due to the sparse firing of our recordings, transfer entropy is biased towards detecting excitatory, rather than inhibitory, interactions. This is because transfer entropy grows with the probability of observing spike events. And, in sparse spike time series it is statistically easier to detect an increase in the number of spikes (an excitatory effect) than it is to detect a decrease in the number of spikes (an inhibitory effect). Thus, here we assume connections are predominantly excitatory.

### Identifying motifs

Synergistic motifs were identified using code inspired by the Matlab Brain Connectivity toolbox [88]. The code was written to categorize all synergistic triads–those in which two transmitters send edges to the same receiver node–according to the set of ten possible synergistic motifs, containing up to four additional edges. Because we were only interested in synergistic motifs, we did not consider the entire set of 3-node motifs. In addition, although motifs 5 and 6 (in this paper) would normally be considered conformationally equivalent, here they are distinct due to the consideration of transmitter and receiver node roles.

### Quantifying synergistic integration

Synergistic integration was measured as synergy. Synergy measures the additional information regarding the future state of the receiver, gained by considering the prior state of the senders jointly, beyond what they offered individually, after accounting for the redundancy between the sending neurons and the past state of the receiver itself. Synergy was calculated according to the partial information decomposition (PID) approach described by [7], including use of the *I_min_* term to calculate redundancy (see Supplemental Material). PID compares the measured bivariate TE between neurons *TE(J→I)* and *TE(K→I)* with the measured multivariate TE (the triad-level information transmission) among neurons *TE({J,K}→I)* to estimate terms that reflect the unique information carried by each neuron, the redundancy between neurons, and the synergy (i.e., gain over the sum of the parts) between neurons. Redundancy was computed as per Supplemental equations 8-10, in which it represents the minimum information (*I_min_*) that J or K provides about each state of I, averaged over all states, and conditioned on the past state of I. Thus, it is the overlapping information—the minimum of that provided by J or K—and can therefore be viewed as redundancy. Synergy was then computed via:

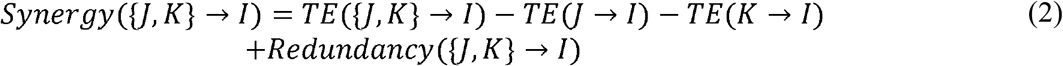

All information terms (synergy, redundancy, and multivariate transfer entropy) were normalized by the entropy of the receiving neuron in order to reflect the proportion of receiver variance for which they accounted and to control for variable entropy across triads and networks. We prefer this normalization to both raw values and other normalizations due to its improved interpretability—rather than raw bits, we analyzed the proportion of receiver variance accounted for. In addition, this particular normalization anchored the analysis with respect to the computing neuron (receiver), which was particularly crucial in the context of this motif-style analysis.

### Statistics

All results are reported as medians or means followed by the 95% bootstrap confidence limits (computed using 10,000 iterations) reported inside of square brackets. Accordingly, figures depict the medians or means with errorbars reflecting the 95% bootstrap confidence limits. Comparisons between conditions or against null models were performed using the nonparametric Wilcoxon signed-rank test, unless specified otherwise. The threshold for significance was set at 0.05, unless indicated otherwise in the text. Bonferroni-Holm corrections were used in cases of multiple comparisons.

## Acknowledgements

We thank Blanca Gutierrez Guzman for helpful comments and discussion.

