## Supporting Information for "Synergistic neural integration is greater downstream of recurrent information flow in organotypic cortical cultures"

SUPPLEMENTAL METHODS

The methods used in this study are described below in the following order: (1) in vitro data collection; (2) construction of effective connectivity networks; and (3) quantification of synergy and related information measures.

***In vitro* data collection**

All procedures were performed in strict accordance with guidelines from the National Institutes of Health, and approved by the Animal Care and Use Committees of Indiana University and the University of California, Santa Cruz.

To study the relationship between neural computation and topological measures of networks of spiking neurons, we analyzed data collected *in vitro*. Data were spontaneously spiking organotypic cultures of mouse somatosensory cortex obtained from postnatal Day 6 to 7 Black 6 mouse pups (RRID:Charles_River:24101632, Harlan) following methods described by [65,89]. Spontaneous (as opposed to stimulus-driven) spiking activity in the cultures was recorded at a high temporal resolution of 50 μs, between 2 and 4 weeks after culture preparation, using a 512-microelectrode array [86]. Array electrodes were flat, 5 μm in diameter and arranged in a triangular lattice with an interelectrode distance of 60 μm. This spacing means that the spiking of most cells is picked up by multiple sites and there are few gaps where cells are too far from electrodes to be recorded. The full array allowed for a total recording area of approximately 0.9 mm by 1.9 mm. This preparation and recording method enabled the isolation of large numbers of neurons (a median of 310 cells per recording in 25 hour-long recordings) at high temporal resolution, beyond what can currently be done in any *in vivo* setup. Crucially, the temporal resolution of this method was small enough to resolve synaptic delays typically found in cortex [43,90].

Once the data were collected, spikes were sorted using a PCA approach based on waveforms detected at seven adjacent electrodes [41,42,86]. This process yielded a single set of spike times for each isolated neuron. Neurons that spiked fewer than 100 spikes during the hour long recording were removed from the analysis. Spike trains were then used to build networks.

**Effective connectivity network construction**

Because neural computation is fundamentally a dynamic process, we focused on examining networks of effective connectivity. In these networks, connections—or edges—represent a predictive relationship between the firing of two different neurons. Note, effective connectivity differs from structural connectivity (synapses or gap junctions between neurons) and functional connectivity (e.g., cross-correlations between neuronal time series). Here, effective connections represent directed information transfer between neurons.

Networks of effective connectivity, representing global activity in recordings, were constructed following methods described previously [8,41] using a measure from information theory known as transfer entropy (TE) [87]. TE was selected for its ability to detect nonlinear interactions and deal with discrete data, such as spike trains. To capture neuron interactions at timescales relevant to synaptic transmission (1-14 ms) [43,44], multiple windows are used to improve the sensitivity to functional interactions across these delays. Spiking data was binned at three logarithmically-spaced bin sizes (1, 1.6 and 3.5 ms) and TE was computed at delays (0-3 bins, for bins of size 1 and 1-4 bins for bins of size 1.6 and 3.5 ms) corresponding to synaptic delays, as in [8,41]. Thus, we computed TE at three timescales, 0.05–3 ms, 1.6–6.4 ms and 3.5–14 ms. Timescales were purposefully designed to be overlapping so that no interactions were neglected. See S1 Fig for an overview of the binning structure used in TE calculations.

TE quantifies an effective connection from neuron J to neuron I by measuring how much information the past state of the neuron J time series ($J_{t-1}$ ) produces regarding the current state of the neuron I time series ($I_{t}$ *)*, beyond what is provided by the past state of the neuron I time series ($I_{t-1}$). Here, time series are binary spike trains for neurons I and J, containing 0 for time bins in which the neuron did not spike and 1 for time bins in which it did spike. Generally, the TE from neuron J to neuron I is computed as:

$${TE}_{J \to I}=\sum_{i_{t},i_{t-1}, j_{t-1}} p\left( i_{t}, i_{t-1}, j_{t-1} \right)\log\left( \frac{p(i_{t}| i_{t-1}, j_{t-1})}{p\left( i_{t} | i_{t-1} \right)} \right)$$

(1)

The probabilities in Eqn. 1 are computed by counting the number of occurrences of all possible combinations of spiking and not spiking in the $i_{t}, i_{t-1}$ and $j_{t-1}$ time bins (of the $I_{t}$, $I_{t-1}$ and $J_{t-1}$ time series) for all bins making up the hour-long recording.

Because we wanted to consider interactions at various timescales associated with synaptic and extra-synaptic transmission, we included a delay between the past and future states of the neurons so that $i_{t-1}$ became $i_{t-d}$ and $j_{t-1}$ became $j_{t-d}$. Additionally, in order to ensure overlapping timescales, we combined the $i_{t-d}$ and $j_{t-d}$ bins with their previous time bins, such that a spike in either or both time bins corresponded to a state of 1 while no spikes in either time bin corresponded to a state of 0 (see S1 Fig for binning structure). Denoting these new bins as $i_{t-d}^{'}$ and $j_{t-d}^{'}$ gives a slightly different form for TE:

$${TE\left( d \right)}_{J \to I}=\sum_{i_{t},i_{t-d}^{'}, j_{t-d}^{'}} p\left( i_{t}, i_{t-d}^{'}, j_{t-d}^{'} \right)\log\left( \frac{p(i_{t}| i_{t-d}^{'}, j_{t-d}^{'})}{p\left( i_{t} | i_{t-d}^{'} \right)} \right)$$

(2)

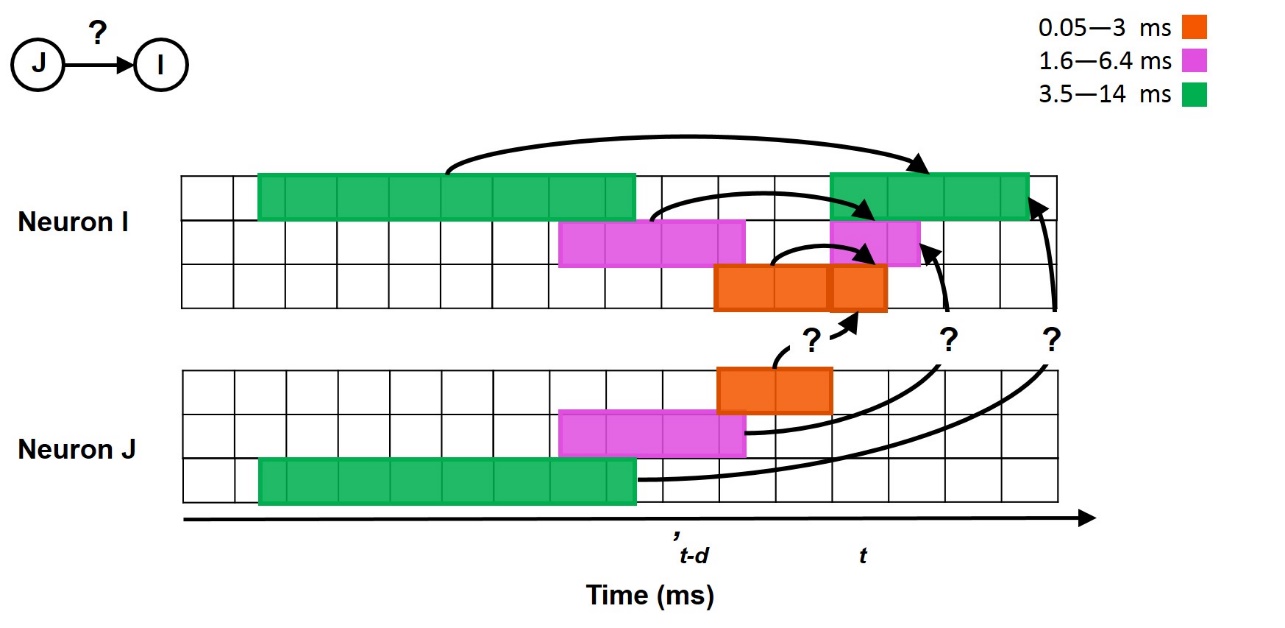

**S1 Fig. Overview of time series binning structure used in transfer entropy calculations.** Transfer entropy was used to quantify a directed, functional connection from neuron J to neuron I which represents how well the current state (t) of neuron I can be predicted by the past state (’t-d) of neuron J, beyond what is known from the past state of neuron I itself. Three synaptic timescales were considered, each with corresponding delays (d). These timescales considered transfer entropy from 0.05­—3 ms, 1.6—6.4 ms, and 3.5—14 ms.

To cast TE in terms of the percentage of the receiver neuron’s capacity that can be accounted for by the transmitting neuron, rather than it representing the amount of information being transmitted from transmitter to receiver, we normalized TE by the entropy of the receiver neuron via:

$${{TE}_{Norm}\left( d \right)}_{J \to I}= \frac{{TE\left( d \right)}_{J \to I}}{-\sum_{i_{t}} p(i_{t})log( p(i_{t}))}$$

(3)

Computing (normalized) TE in this way between all pairs of binned neuronal time series results in a time-scale dependent, weighted, directed network. Networks are weighted because some pairs of neurons fire more frequently and reliably at certain delays than others, and they are directed because a predictive, statistical relationship that exists from neuron J to neuron I, may not exist from neuron I to neuron J. Each element $a_{ij}$ in the TE matrix is the TE value from the $i^{th}$ to the $j^{th}$ neuron. TE values of zero denote the absence of an effective connection between the two neurons, while TE values greater than zero represent the weighted strength of the effective connection between the two neurons.

To determine the significance of network edges (TE values), TE values were computed for 5000 pairs of jittered spike trains. Spike trains were jittered by randomly adjusting the timing of each spike by a small amount proportional to the timescale being examined. This preserved the overall firing of each neuron, as well as the longer timescale dynamics, but destroyed the precise spike timing between neurons. Only spike trains of source neurons were jittered in order to preserve the auto-prediction in the target neuron. Spike trains were jittered according to a uniform distribution with a width of seven bins centered on the observed location of each spike. This procedure is discussed in more detail in [41]. TE values which were larger than 99.9% of jittered values were considered significant, corresponding to an alpha value of 0.001. Computing significant, normalized TE values for 25 recordings at three timescales resulted in 75 full networks.

An important consideration in the calculation of our TE values, which includes a delay in the past of the receiver, is that we may be overestimating TE and increasing our odds for detecting false positives [91]. Importantly, we control for the increased risk of false-positives by using an alpha-level of 0.001. However, the potential overestimation of TE comes from more variance being attributed to the sender neurons than to the receiver neuron itself. Despite the advantages of the [91] method, which does not include a delay in the receiver past, we chose the current method in order to isolate interactions at separate timescales.

To ensure that the detection of motifs was not biased by the spatial sampling of the recording apparatus, we compared the distances between nodes in each motif to the distances between all nodes, across networks. To do this, we determined the distribution of node distances for each motif, combined across networks to provide enough samples of each motif type. We then compared these 10 distributions to the overall distribution of distances between nodes. We found no significant differences between distributions of motif neuron distances and the overall distribution of distances, excepting in the case of motif 10 (KS tests revealed that distance distributions for motifs 1-9 were not significantly different from the overall distance distribution at the α = 0.005 level; S2 Fig). Distances between nodes in motif 10 are different from the overall distances (mean +/- s.d.: 0.57 mm [0.13 mm, 1.01 mm] versus 0.74 mm [0.33 mm, 1.16 mm]) because considerably fewer nodes exhibit motif 10 (the rarest motif). Thus, there are fewer nodes to span the full recording space (S3 Fig).

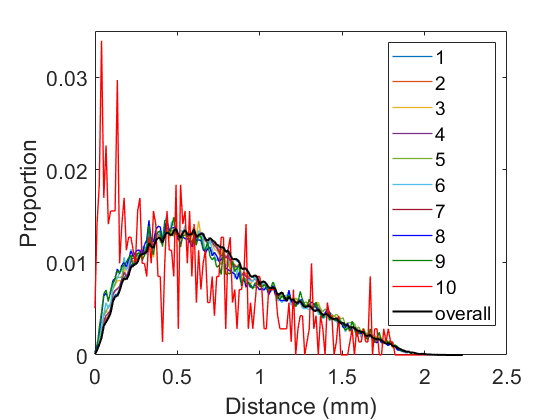

**S2 Fig. Distributions of node distances for motifs 1-9 are not significantly different from the overall distribution of distances.** Distribution of distances for motif 10 is significantly different from the overall distribution of distances, due to a greater prevalence of smaller distances.

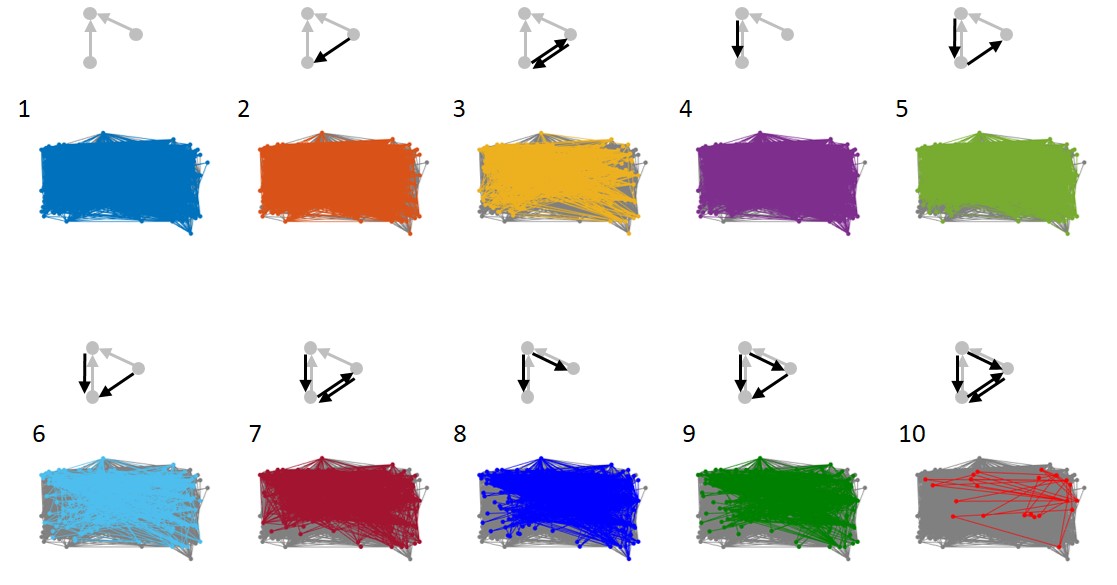

**S3 Fig. Spatial distribution of motifs 1-9 is not significantly different from the spatial distribution of the rest of the network.** The spatial distribution—on the recording array­—of motifs (colored nodes and edges) relative to the rest of the network (gray nodes and edges) is shown for a representative network. The spatial distribution covers a smaller range for motif 10, of which there are relatively few cases.

To be confident that our timescales captured the peak of information processing in our networks, we calculated TE at delays other those analyzed here for two representative networks. First, spike trains were binned at 1 ms. Then TE was calculated at multiple delays ranging from 0 to 501 ms, in steps of 3 ms, for all existing pairs of significant effective connections in the network. We found that TE tended to peak in the 1-14 ms delay range for most effective connections (S4 Fig). Across networks, 87.3% of pairs, on average, had a peak of TE between 1 and 14 ms.

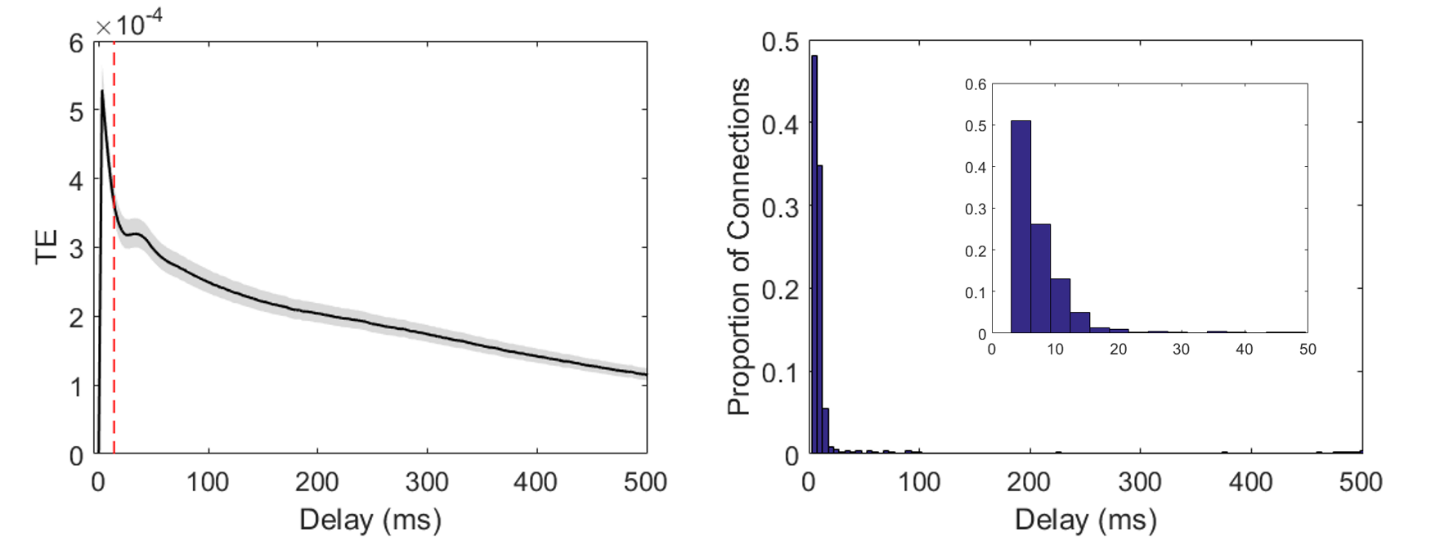

**S4 Fig. TE peaks between 1-14 ms.** Mean distribution of TE over time for all effective connections from two representative networks. Left: The black line shows the mean TE over all effective connections from two representative networks. The shaded region shows the 95% confidence interval. The vertical dashed red line indicates the upper bound of the synaptic timescales. Across connections, the peak TE occurs below this bound at short latencies. Right: Histogram of the delay to the maximum TE over connections. The height of each bar shows the proportion of connections for which the peak TE was found to occur at the delay indicated along the x-axis. Most connections had max TE at short delays as shown in the inset panel which zooms in to the first 50 ms of the x-axis. These plots show that most connections had a peak TE at less than 14 ms.

**Quantification of synergy**

Computation by neurons receiving inputs from two other neurons in these networks was quantified following the partial information decomposition (PID) method from [7]. The PID allows multivariate TE (mvTE) to be separated into distinct information components, one of which is a measure of neural computation termed synergy. The general form of the decomposition of multivariate TE between three neuronal time series, with two transmitter neurons, J and K, each sending a single input to one receiver neuron, I, can be expressed as:

$$TE\left( \left\{ J,K \right\}\to I \right)= \mathrm{Synergy}\left( \left\{ J,K \right\}\to I \right)+ \mathrm{Unique}\left( K;J\to I \right)+$$

$$\mathrm{Unique}\left( J;K\to I \right)+ \mathrm{Redundancy}(\{J,K\}\to I)$$

(4)

where $\left\{ J,K \right\}$ is a vector of the combined J and K time series (S5 Fig). Similarly, we can express the decomposition of bivariate TE from neuron J to I and neuron K to I as:

| $TE\left( J\to I \right)= \mathrm{Unique}\left( K;J\to I \right)+ \mathrm{Redundancy}(\{J,K\}\to I)$ | (5) |
| --- | --- |
| and |  |
| $TE\left( K\to I \right)= \mathrm{Unique}\left( J;K\to I \right)+ \mathrm{Redundancy}(\{J,K\}\to I)$ | (6) |

In Equations 4-6, all terms are quantified in units of bits (see [7] for a full description of these terms). The unique terms correspond to the information provided by that time series alone (either the J or the K time series) about the current state of I. The redundant term represents the overlapping information provided by time series J and K about the current state of I. Notice, in Equations 5 and 6, that although TE is only dependent on the two time series that are directly interacting (either J and I, or K and I), because J and K are both interacting with the same time series, their unique interactions are influenced by each other. Thus, the unique information provided by one of these time series is dependent on the other. In other words, because J and K provide some redundant (overlapping) information about I, J influences how much information K provides uniquely versus redundantly about I. Likewise, K influences how much information J provides uniquely versus redundantly about I.

The synergistic term in Equation 4 is the additional information (beyond the unique and redundant information) that is accounted for in the receiver activity (I) based on the non-overlapping information from both inputs (J and K) occurring simultaneously. Thus, synergy is a proxy for the non-linear computation which takes information from two sources and combines them in some way to generate a unique output. Examples of high synergy computations include ‘AND’ and ‘XOR,’ wherein knowledge of the state of both upstream neurons is needed to predict the state of the receiving neuron [8].

To calculate synergy, note that Equation 4 can be rewritten as:

$$\mathrm{Synergy}\left( \left\{ J,K \right\}\to I \right)= TE\left( \left\{ J,K \right\}\to I \right)- TE\left( J\to I \right)-$$

$$TE\left( K\to I \right)+ \mathrm{Redundancy}(\{J,K\}\to I)$$

(7)

by substituting Equations 5 and 6 and solving for synergy. Notice that we can compute all TE terms in Equation 7 via Equation 1. This leaves only the Redundancy term to be computed. We use the method for estimating redundancy provided by [7]. That is redundancy is defined as follows in terms of a quantity titled the minimum information $I_{\min}$:

$$\mathrm{Redundancy}\left( \left\{ J,K \right\}\to I \right)≝I_{\min}\left( I_{t};J_{t-1}K_{t-1}|I_{t-1} \right)=$$

$$\sum_{i_{t}} p\left( i_{t} \right)\min_{R\in\left\{ J_{t-1},K_{t-1} \right\}} I_{spec}\left( I_{t}= i_{t};R | I_{t-1} \right)=$$

$$\sum_{i_{t}} p\left( i_{t} \right)\min_{R\in\left\{ J_{t-1},K_{t-1} \right\}} [I_{spec}\left( I_{t}=i_{t};R,I_{t-1} \right)-I_{spec}\left( I_{t}=i_{t};I_{t-1} \right)]$$

(8)

where the specific information $I_{spec}$ is defined as:

$$I_{spec}\left( I_{t}=i_{t};R,I_{t-1} \right)=\sum_{r, i_{t-1}} p\left( r,i_{t-1}|i_{t} \right)\log\left( \frac{p(r, i_{t-1}, i_{t})}{p\left( r,i_{t-1} \right)p(i_{t})} \right)$$

(9)

and

$$I_{spec}\left( I_{t}=i_{t};I_{t-1} \right)=\sum_{i_{t-1}} p\left( i_{t-1}|i_{t} \right)\log\left( \frac{p(i_{t-1}, i_{t})}{p\left( i_{t-1} \right)p(i_{t})} \right)$$

(10)

Thus, redundancy is the minimum information provided by J or K about each state of I, averaged over all possible states. In other words, redundancy is the overlapping information (the shared information) that the past states of J and K provide about the current states of I. Redundancy was calculated via Equations 9 and 10. Finally, synergy was calculated via Equation 7. Computing synergy for all possible triads (for each neuron that received at least two significant inputs, all possible groupings of two input neurons and the receiver were considered) in all networks yields a single synergy value per triad. We then normalized synergy (as well as redundancy and mvTE) values by dividing by the entropy of the future state of I, as done in Equation 3.

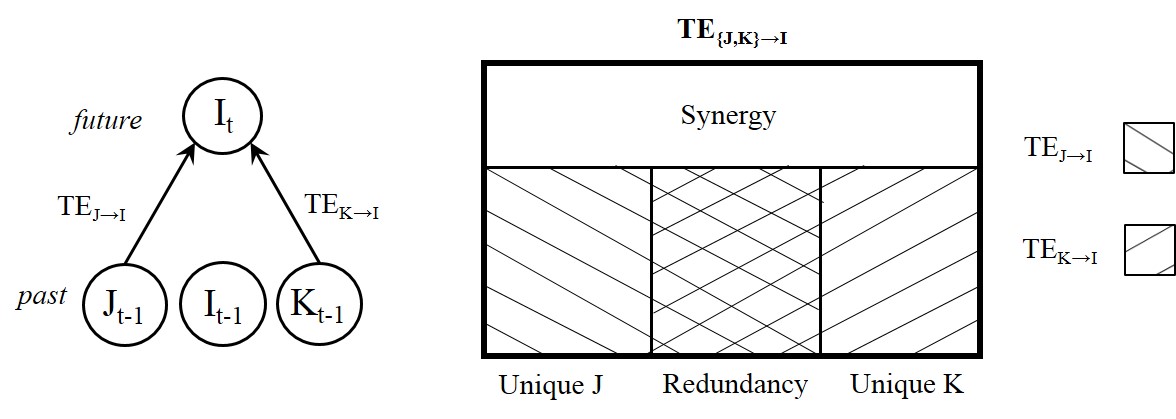

**S5 Fig. The Partial Information Decomposition.** In this study, we analyzed two-input computations which were determined using the Partial Information Decomposition to dissect multivariate transfer entropy (occurring among three neurons, with two transmitter neurons each sending significant information to a receiver neuron) into synergistic, redundant, and unique information terms. The synergistic information component was used to represent the amount of computation carried out by the receiver.

SUPPLEMENTAL RESULTS

Below are additional results not presented in the main text pertaining to: (1) alternative normalizations of synergy; (2) relationships between associated information terms­—redundancy and multivariate transfer entropy—and recurrence and feedback; (3) an alternative implementation of synergy; and (4) synergy results at separate timescales. All figures in this section are plotted as in Fig 4 of the main text. Tables accompanying figures show results of the repeated measures ANOVA, including main effects and interaction effects, as well as correlation coefficients. Tables are arranged as in Table 1 of the main text.

**Alternative normalizations of synergy**

The current study showed results of raw synergy and synergy normalized by receiver entropy. Here, we show additional normalizations of synergy by the multivariate transfer entropy (Syn/mvTE; S6 Fig) and by the feedforward edge weight (Syn/FF; S7 Fig). As with both raw synergy and synergy normalized by receiver entropy, we observed that synergy normalized by mvTE and by FF increased significantly with the number of recurrent edges.

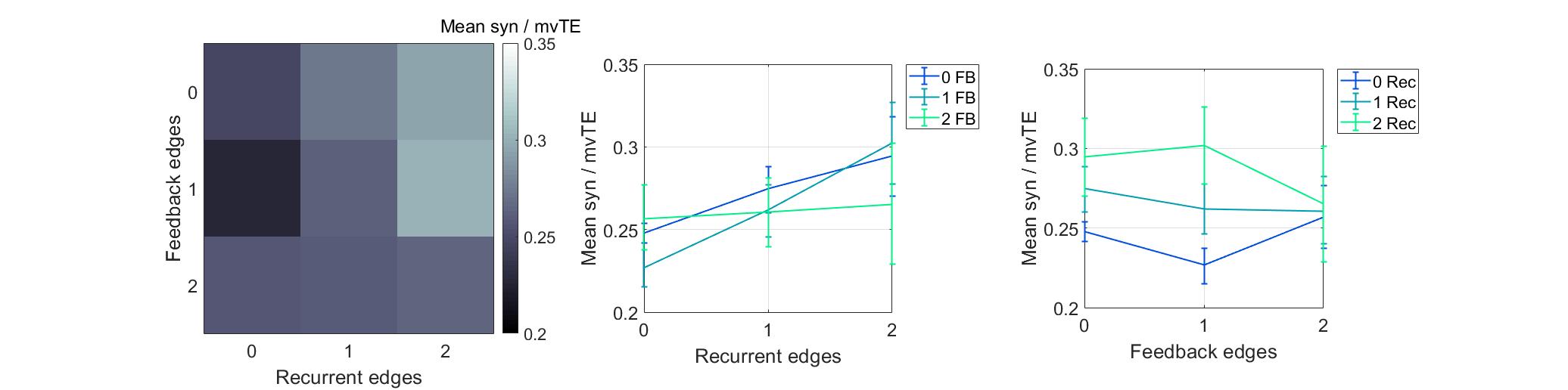

| **Syn / mvTE** | df | F | p_ANOVA_ | rho | p_rho_ |
| --- | --- | --- | --- | --- | --- |
| Recurrent | 2, 148 | 14.4 | **<1x10^-5^** | 0.29 | **<1x10^-10^** |
| Feedback | 2, 148 | 35.8 | **<1x10^-12^** | -0.08 | 0.06 |
| Recurrent x Feedback | 4, 296 | 11.3 | **<1x10^-7^** | -- | -- |

**S6 Fig. Synergy normalized by multivariate transfer entropy increased with the number of recurrent edges.** (Left) Motifs are ordered based on the number of recurrent edges (columns) and feedback edges (rows). The heatmap depicts brighter colors where there are larger normalized synergy values. (Middle) Curves representing rows in (Left), plotted with errorbars computed across networks, show that synergy increased as the number of recurrent edges increased. (Right) Curves representing columns shown in (Left), plotted with errorbars computed across networks, show that synergy decreased as the number of feedback edges increased. Errorbars are 95% bootstrap confidence intervals around the mean. **Table**. Relationships between normalized synergy and recurrence and feedback. Columns 1-3 (*df, F, p_ANOVA_*) show the results of a repeated measures ANOVA for the normalized synergy predicted by the number of recurrent and feedback edges. Columns 4-5 (*rho, p_rho_*) show the results of Spearman rank correlations between normalized synergy and the number of recurrent and feedback edges. P-values significant at the α=0.05 level are in bolded font.

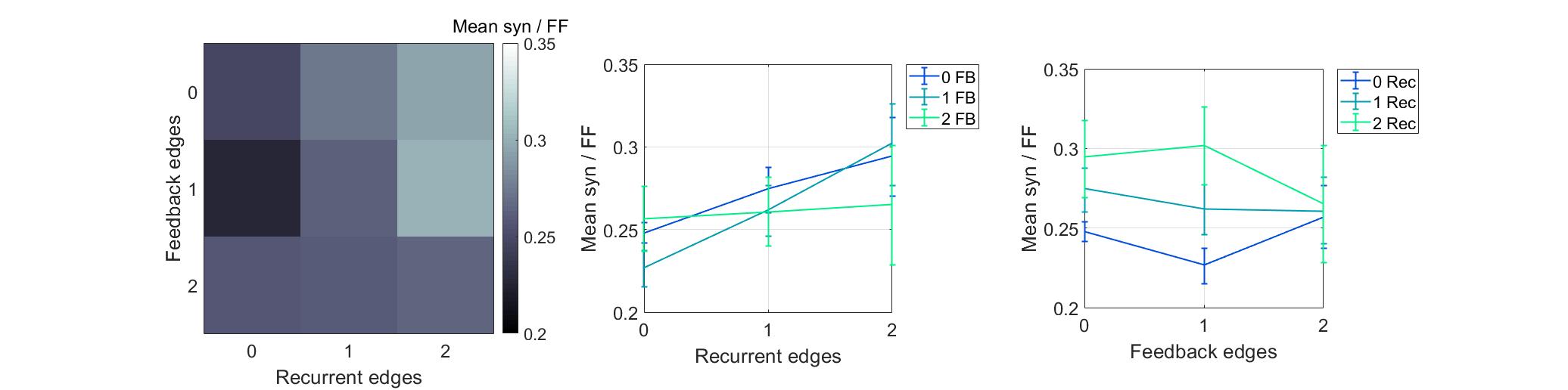

| **Syn / FF** | df | F | p_ANOVA_ | rho | p_rho_ |
| --- | --- | --- | --- | --- | --- |
| Recurrent | 2, 148 | 13.4 | **<1x10^-5^** | 0.28 | **<1x10^-10^** |
| Feedback | 2, 148 | 32.3 | **<1x10^-11^** | -0.06 | 0.17 |
| Recurrent x Feedback | 4, 296 | 10.9 | **<1x10^-7^** | -- | -- |

**S7 Fig. Synergy normalized by the feedforward edge weight increased with the number of recurrent edges.** Results plotted as in S6 Fig.

**Redundancy and MvTE**

In addition to synergy, another important information term obtained from partial information decomposition of the multivariate transfer entropy is redundancy, which measures the amount of shared information the sender neurons provide about the receiver. To offer context to the relationships studied here, below we include the results of relating both redundancy and the multivariate transfer entropy to the number of recurrent and feedback edges in motifs. As in the manuscript, results are shown for information terms normalized by receiver entropy, as well as for raw information terms. We also include redundancy normalized by feedforward edge weight, which is nearly identical to normalization by multivariate transfer entropy (mvTE), to compare to additional synergy normalizations above. As with synergy, we observed that both redundancy (red)—whether normalized by receiver entropy (S8 Fig), raw (S10 Fig), or normalized by feedforward edge weight (S12 Fig)—and mvTE—whether normalized by receiver entropy (S9 Fig) or raw (S11 Fig)—increase significantly with the number of recurrent edges. These results agree with previous findings that normalized synergy is positively correlated with both redundancy and multivariate transfer entropy at synaptic timescales [10].

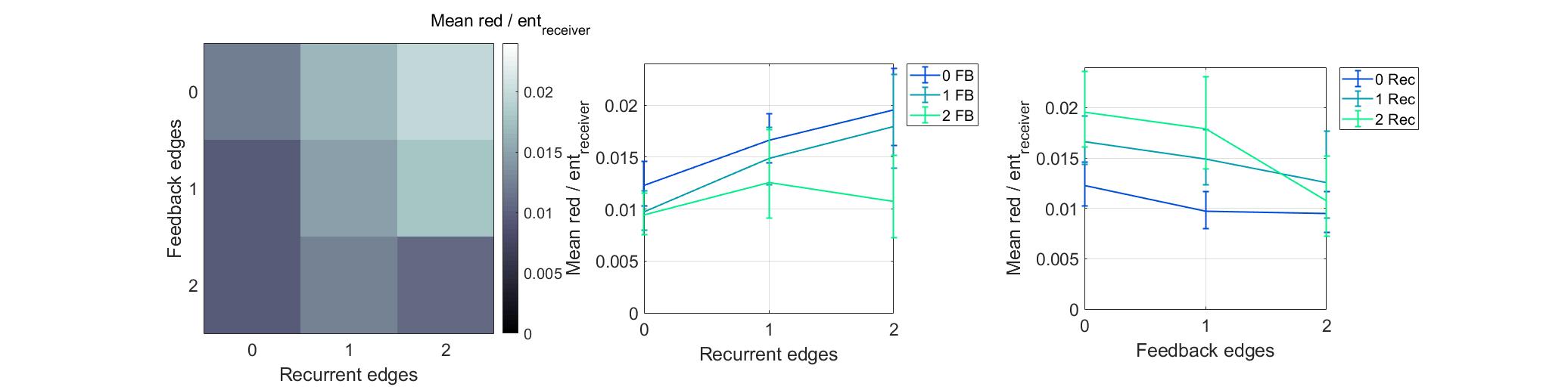

| **Red / ent_receiver_** | df | F | p_ANOVA_ | rho | p_rho_ |
| --- | --- | --- | --- | --- | --- |
| Recurrent | 2, 148 | 6.7 | **<0.01** | 0.20 | **<1x10^-5^** |
| Feedback | 2, 148 | 51.3 | **<1x10^-17^** | -0.23 | **<1x10^-7^** |
| Recurrent x Feedback | 4, 296 | 5.2 | **<1x10^-3^** | -- | -- |

**S8 Fig. Redundancy normalized by receiver entropy increased with the number of recurrent edges.** Results plotted as in S6 Fig.

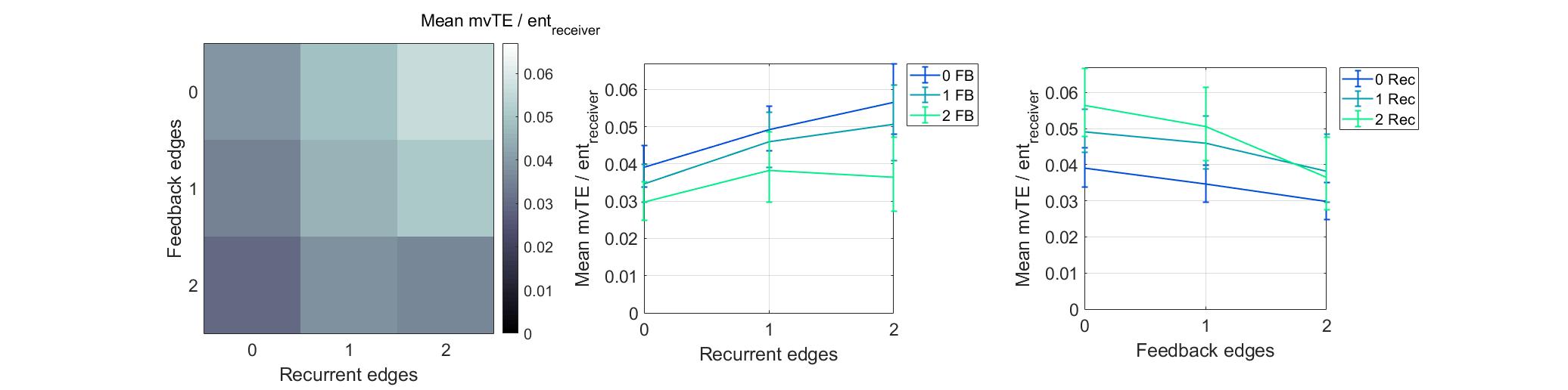

| **MvTE / ent_receiver_** | df | F | p_ANOVA_ | rho | p_rho_ |
| --- | --- | --- | --- | --- | --- |
| Recurrent | 2, 148 | 8.8 | **<1x10^-3^** | 0.19 | **<1x10^-5^** |
| Feedback | 2, 148 | 65.9 | **<1x10^-17^** | -0.21 | **<1x10^-6^** |
| Recurrent x Feedback | 4, 296 | 4.4 | **<0.01** | -- | -- |

**S9 Fig. Multivariate transfer entropy normalized by receiver entropy increased with the number of recurrent edges.** Results plotted as in S6 Fig.

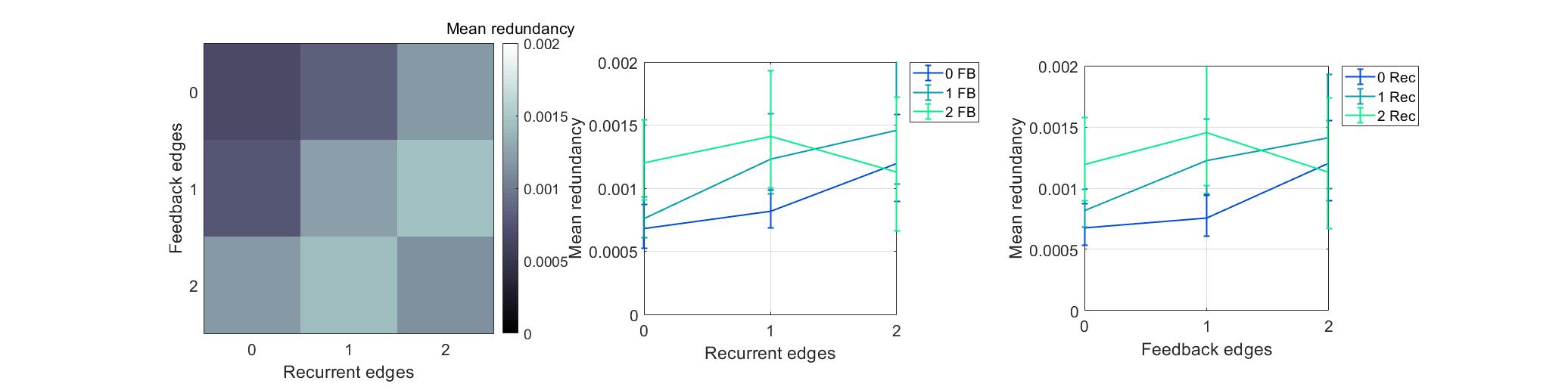

| **Redundancy** | df | F | p_ANOVA_ | rho | p_rho_ |
| --- | --- | --- | --- | --- | --- |
| Recurrent | 2, 148 | 3.8 | **0.03** | 0.12 | **<0.01** |
| Feedback | 2, 148 | 3.3 | **0.04** | 0.07 | 0.1 |
| Recurrent x Feedback | 4, 296 | 7.2 | **<1x10^-4^** | -- | -- |

**S10 Fig. Raw redundancy increased with the number of recurrent edges.** Results plotted as in S6 Fig.

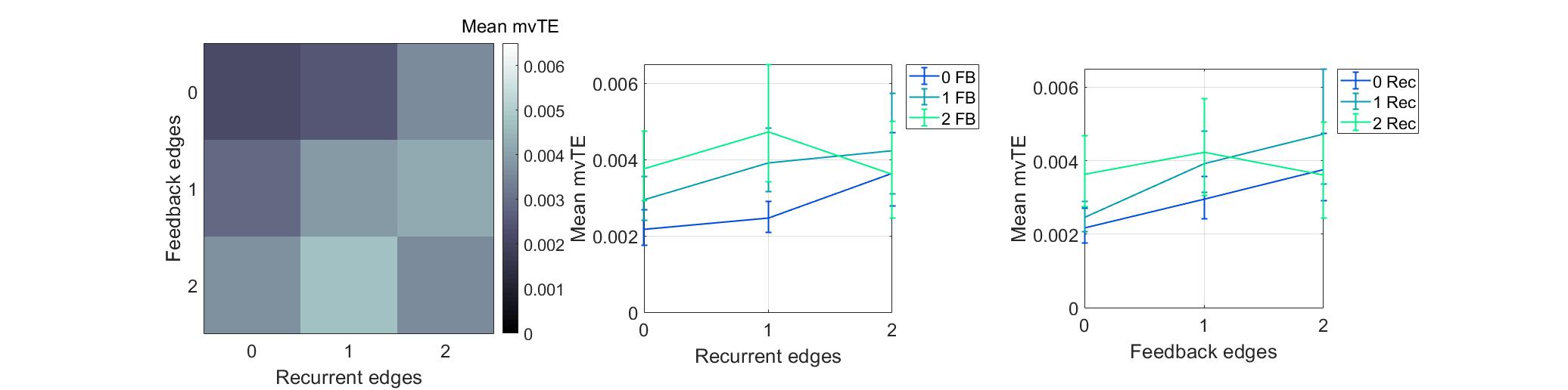

| **MvTE** | df | F | p_ANOVA_ | rho | p_rho_ |
| --- | --- | --- | --- | --- | --- |
| Recurrent | 2, 148 | 7.1 | **<0.01** | 0.09 | **0.04** |
| Feedback | 2, 148 | 4.9 | **<0.01** | 0.13 | **<0.01** |
| Recurrent x Feedback | 4, 296 | 7.5 | **<1x10^-5^** | -- | -- |

**S11 Fig. Raw multivariate transfer entropy increased with the number of recurrent edges.** Results plotted as in S6 Fig.

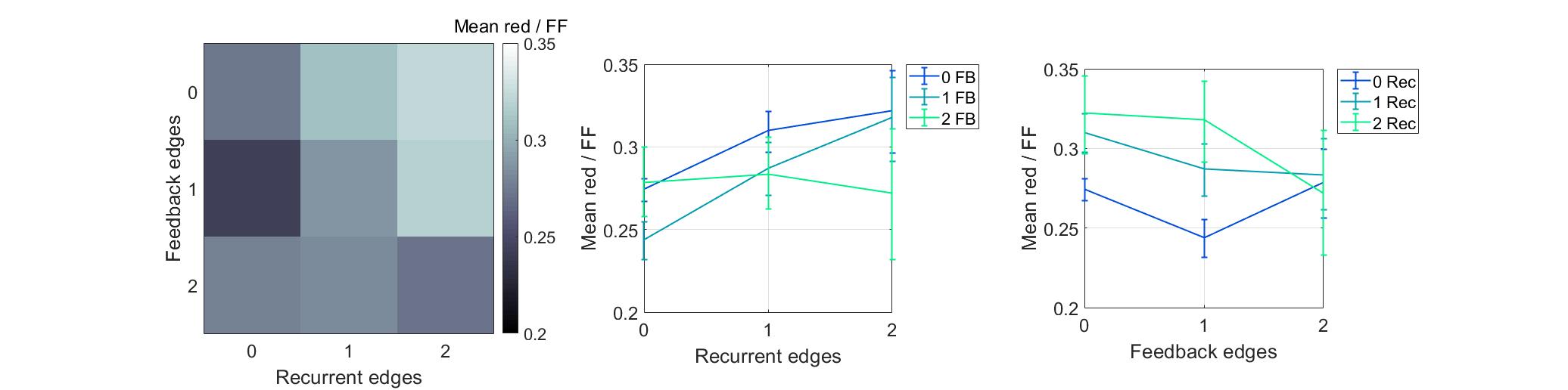

| **Redundancy / FF** | df | F | p_ANOVA_ | rho | p_rho_ |
| --- | --- | --- | --- | --- | --- |
| Recurrent | 2, 148 | 19.3 | **<1x10^-7^** | 0.28 | **<1x10^-10^** |
| Feedback | 2, 148 | 47.2 | **<1x10^-15^** | -0.13 | **<0.01** |
| Recurrent x Feedback | 4, 296 | 11.9 | **<1x10^-8^** | -- | -- |

**S12 Fig. Redundancy normalized by the strength of feedforward edges increased with the number of recurrent edges.** Results plotted as in S6 Fig.

**Alternative implementation of synergy**

To ensure that our findings were not dependent on the method used to calculate synergy, we performed additional analyses which implemented an alternative method. In this alternative method, we considered the effect of calculating the lower bound on synergy, which we refer to as “bonafide” synergy. This method also uses PID but sets redundancy to be the smallest possible value. Effectively, in this approach synergy is computed as follows:

$$\mathrm{Synergy}\left( \left\{ J,K \right\}\to I \right)= \mathrm{argMax}[ TE(\{J,K\}\to I) - TE\left( J\to I \right) - TE\left( K\to I \right), 0 ]$$

(11)

Consequently, synergy is minimized or set to zero when the sum of $TE(J\to I)$ and $TE(K\to I$) is greater than$TE(\{J,K\}\to I)$. Note, $TE(\{J,K\}\to I) - TE\left( J\to I \right) - TE\left( K\to I \right)$ is equivalent to $\mathrm{Synergy}\left( \left\{ J,K \right\}\to I \right)-\mathrm{Redundancy}\left( \left\{ J,K \right\}\to I \right).$

When we used these synergy values and repeated our core analysis, we observed that bonafide synergy—normalized by receiver entropy (S13 Fig) and raw (S14 Fig)—increased with the number of recurrent edges.

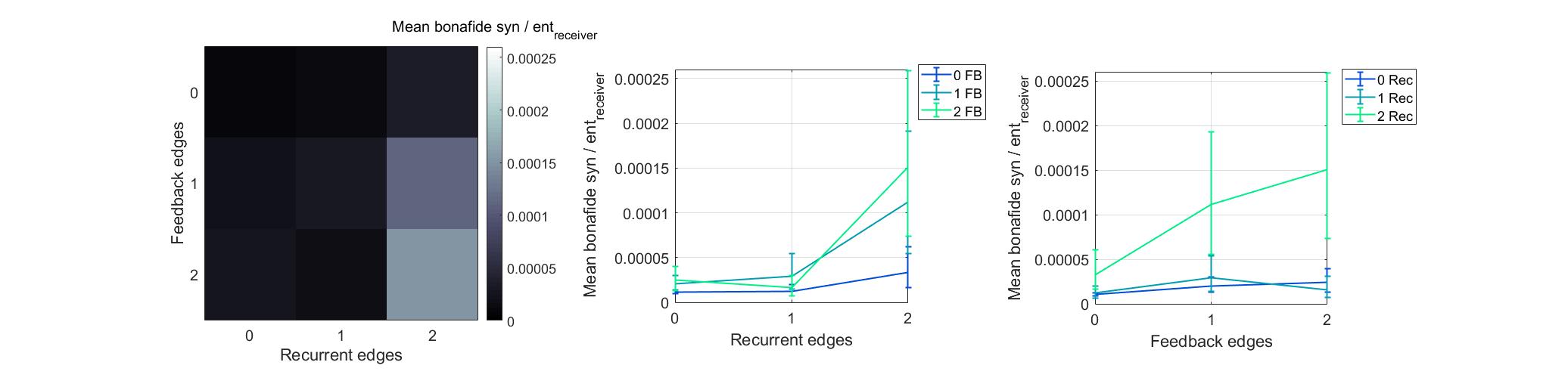

| **Bonafide Syn / ent_receiver_** | df | F | p_ANOVA_ | rho | p_rho_ |
| --- | --- | --- | --- | --- | --- |
| Recurrent | 2, 148 | 7.6 | **<1x10^-3^** | -0.009 | 0.83 |
| Feedback | 2, 148 | 4.3 | **0.02** | -0.03 | 0.43 |
| Recurrent x Feedback | 4, 296 | 1.7 | 0.15 | -- | -- |

**S13 Fig. Bonafide synergy normalized by receiver entropy increased with the number of recurrent edges.** Results plotted as in S6 Fig.

The relationship between normalized bonafide synergy and number of feedback edges is positive, unlike the relationship between normalized synergy and number of feedback edges, which is negative. This is likely due to the fact that the bonafide analysis sets a high threshold for synergy. Thus, the triads that are recruited into the analysis are those with generally high synergy, and therefore haven’t been negatively affected by the presence of feedback.

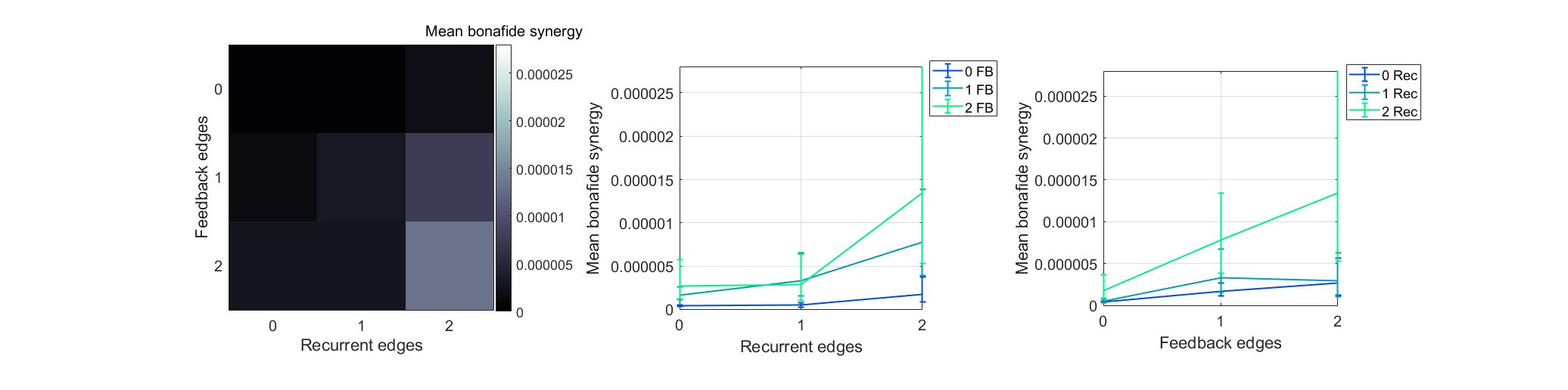

| **Bonafide Synergy** | df | F | p_ANOVA_ | rho | p_rho_ |
| --- | --- | --- | --- | --- | --- |
| Recurrent | 2, 148 | 4.1 | **0.02** | -0.03 | 0.54 |
| Feedback | 2, 148 | 4.8 | **0.01** | 0.08 | **0.05** |
| Recurrent x Feedback | 4, 296 | 0.7 | 0.61 | -- | -- |

**S14 Fig. Raw bonafide synergy increased with the number of recurrent edges.** Results plotted as in S6 Fig.

**Synergy at separate timescales**

The current study investigated the relationship between synergy and number of recurrent and feedback edges across three overlapping timescales relevant to synaptic connectivity. The results were pooled in the manuscript. Here we show them for each timescale separately. The pattern of results observed when pooled held when examined separately at each timescale (S15 Fig).

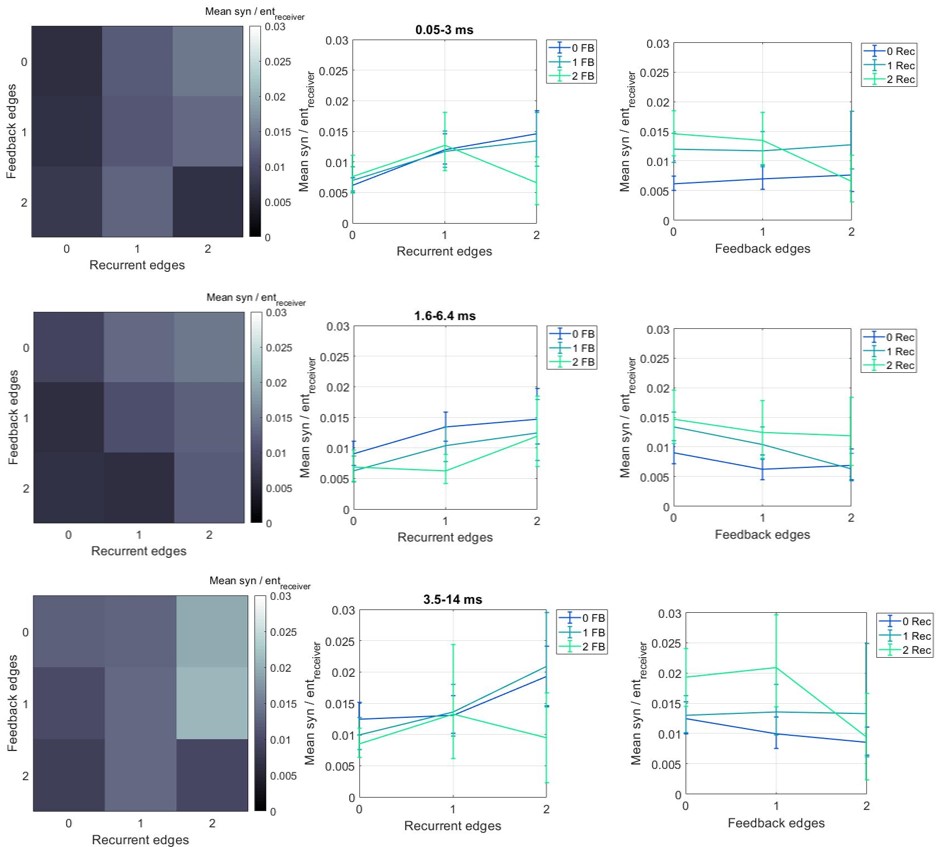

| **0.05—3 ms** | df | F | p_ANOVA_ | rho | p_rho_ |
| --- | --- | --- | --- | --- | --- |
| Recurrent | 2, 48 | 13.2 | **<1x10^-4^** | 0.34 | **<1x10^-5^** |
| Feedback | 2, 48 | 16 | **<1x10^-5^** | -0.09 | 0.26 |
| Recurrent x Feedback | 4, 96 | 4.8 | **<0.01** | -- | -- |
| **1.6—6.4 ms** | *df* | *F* | *p*_ANOVA_ | *rho* | *p_rho_* |
| Recurrent | 2, 48 | 4.6 | **0.01** | 0.22 | **<0.01** |
| Feedback | 2, 48 | 22.4 | **<1x10^-6^** | -0.29 | **<1x10^-4^** |
| Recurrent x Feedback | 4, 96 | 3.1 | **0.02** | -- | -- |
| **3.5—14 ms** | *df* | *F* | *p*_ANOVA_ | *rho* | *p_rho_* |
| Recurrent | 2, 48 | 0.7 | 0.5 | 0.24 | **<0.01** |
| Feedback | 2, 48 | 17.7 | **<1x10^-5^** | -0.25 | **<0.001** |
| Recurrent x Feedback | 4, 96 | 2.7 | **0.03** | -- | -- |

**S15 Fig. Synergy normalized by receiver entropy increases with the number of recurrent edges at each timescale separately.** Results plotted as in S6 Fig.

Despite there being no significant main effect of recurrence at the longest timescale (3.5-14 ms), normalized synergy was significantly positively correlated with the number of recurrent edges (Spearman *r* = 0.24, n=225, p <0.01) at the longest timescale.
